# Mechanosensitive control of mitotic entry

**DOI:** 10.1101/2021.09.23.461473

**Authors:** Margarida Dantas, Andreia Oliveira, Paulo Aguiar, Helder Maiato, Jorge G. Ferreira

## Abstract

As cells prepare to divide, they must ensure that enough space is available to assemble the mitotic machinery without perturbing tissue homeostasis^1^. To do so, cells undergo a series of biochemical reactions regulated by cyclin B1-CDK1 that trigger the reorganization of the actomyosin cytoskeleton^2^ and ensure the coordination of cytoplasmic and nuclear events^3^.

Along with the biochemical events that control mitotic entry, mechanical forces have recently emerged as important players in the regulation of cell cycle events^4–6^. However, the exact link between mechanical forces and the biochemical pathways that control mitotic progression remains to be established. Here, we identify a mechanical signal on the nucleus that helps set the time for nuclear envelope permeabilization (NEP) and mitotic entry. This signal relies on actomyosin contractility, which leads to nuclear unfolding during the G2-M transition, activating the stretch-sensitive cPLA2 on the nuclear envelope. This contributes to the spatiotemporal translocation of cyclin B1 to the nucleus. Our data demonstrate how nuclear mechanics during the G2-M transition contribute to timely and efficient mitotic spindle assembly and prevents chromosomal instability.

## Introduction

Cell cycle progression is regulated by cyclins and their associated kinases. One such complex, composed of cyclin B1-CDK1, is responsible for regulating entry into mitosis. The biochemical mechanisms that regulate mitotic entry have been extensively studied in the past (for review see^7^). For most of the cell cycle, the cyclin B1-CDK1 complex is inactive, due to low cyclin B1 expression levels and its mainly cytoplasmic localization^8,9^. As cells approach late G2/prophase of mitosis, cyclin B1 expression increases^10^, resulting in its binding to CDK1. This cyclin B1-CDK1 complex is kept in an inactive state due to Wee1-mediated phosphorylations of CDK1 on residues T14 and Y15^7^. Through the action of Cdc25 phosphatases, these inhibitory phosphorylations are removed, leading to complex activation. The active cyclin B1-CDK1 complex then stimulates its own nuclear import^3^ through the nuclear pore complexes (NPCs), resulting in chromosome condensation^11^ and nuclear envelope permeabilization (NEP)^4^. Consequently, the cyclin B1-CDK1 complex has been proposed to effectively synchronize cytoplasmic and nuclear events^3^, crucial for mitotic entry and efficient spindle assembly.

The influence of mechanical forces on the cell cycle and some of its key regulators has received considerable attention over the last few years^1,6,12–16^. Experiments performed in isolated cells and epithelial layers have demonstrated that mechanical forces can stimulate the G1-S transition^6,12,14,16,17^ by controlling specific transcriptional programs^12,14,16^. This is likely due to forces imposed on the nucleus^18,19^ that induce its flattening^16,20,21^, facilitating the nuclear accumulation of transcription factors (TFs)^20,22^, changing the organization of both chromatin^23^ and the nuclear envelope (NE)^24^ or altering cell contractility^21,25^.

Evidence for mechanical regulation during other cell cycle phases is more limited. Recently, mechanical stretching was proposed to trigger the G2-M transition by activating Piezo 1^13^. Moreover, we and others have shown that cell traction forces decrease during the G2-M transition ^6,17,26^, to allow mitotic cell rounding and efficient cell division^1,26^. Although these findings highlight the interplay between physical forces and cell proliferation, it remains unknown whether the main biochemical events that occur during the G2-M transition are sensitive to mechanical cues. Importantly, whether the spatial and temporal behaviour of the cyclin B1-CDK1 complex responds to mechanical forces and contributes to ensure timely and efficient cell division remains unknown. Here, we investigate whether and how cyclin B1 responds to physical forces during the G2-M transition. We show that nuclear deformation triggers a contractility-mediated mechanism that facilitates the translocation of cyclin B1 to the nucleus, setting the timing of mitotic entry.

## Results and discussion

We started by investigating whether the G2-M transition is sensitive to external stimuli. We seeded RPE-1 cells expressing H2B-GFP/tubulin-RFP in non-adherent, hydrophobic conditions (PLL-g-PEG) and imaged them as they progressed from G2 to mitosis. Under these conditions, cells failed to enter mitosis (Fig. 1a, c), confirming that the G2-M transition requires contact with external stimuli. We then tested whether mechanically stimulating cells in nonadherent conditions was sufficient to stimulate mitotic entry. For this purpose, we used a dynamic cell confinement device^27^, which gives us temporal control over the process. Confinement was achieved by imposing on cells a fixed height of 8μm with a rigid polydimethylsiloxane (PDMS)-coated glass slide (Sup. Fig. 1a). Upon acute mechanical confinement, cells re-gained the ability to enter mitosis (Fig. 1a-c; ***p<0.001), indicating that mechanical confinement is sufficient to overcome the lack of external stimuli. Under confinement conditions, cells in late G2 entered mitosis within approximately 140±80sec (mean±sd) after stimulation, ruling out that this event was due to increased cyclin B1 transcription, as previously described^13^. One alternative hypothesis is that physical confinement accelerates mitotic entry by inducing a premature transport of cyclin B1 to the nucleus, as previously proposed for YAP or MyoD^20,22^. To confirm this is the case, we monitored the dynamics of nuclear accumulation of endogenous cyclin B1 tagged with Venus in RPE-1 cells, in normal and confined conditions (Fig. 1d, e). We defined time zero as the lowest fluorescence intensity levels of nuclear cyclin B1 and quantified its increase as cyclin B1 translocated to the nucleus, up until the moment of tubulin entry. This last event marks the loss of nuclear barrier function, which we defined as NEP. By analysing the pattern of nuclear cyclin B1 translocation in unmanipulated cells, we determined the translocation to occur within 478±102 sec, with a half-time of 331 sec. Strikingly, we verified that mechanical stimulation triggered a fast nuclear accumulation of cyclin B1 (Fig. 1f-h; ***p<0.001). In confined cells, translocation occurred within 101±12 sec, with a half-time of 70sec, which resulted in a faster mitotic entry (Fig. 1f, h). This was not due to a rupture of the nucleus, as we could not detect any obvious tears or gaps in the NE when RPE-1 cells expressing Lap2β-mRFP were confined (Sup. Fig. 1b). Together with the observed delay between cyclin B1 accumulation and tubulin translocation to the nucleus (Fig. 1d, e, h), these findings strongly suggest that the nuclear barrier function remains intact after confinement. Instead, as previously reported^21^, confinement promoted an unfolding of the NE which could be readily observed (Sup. Fig. 1b) and resulted in an increase in the distance between neighbouring NPCs (Sup. Fig. 1c, d; ***p<0,001), when compared to unconfined cells. Importantly, this unfolding was also observed in unmanipulated cells as they progress from interphase to prophase (Sup. Fig. 1d; *** p<0.001). Our data suggests that the mechanical environment might affect cyclin B1 nucleoplasmic translocation. To test this, we seeded cells on a soft hydrogel (5kPa) or on a rigid glass, inducing low or high cellular tension, respectively. As predicted, cells on glass were more efficient in cyclin B1 nuclear shuttling, than cells on a soft gel (Fig. 1i, j***p<0.001). Because DNA damage can delay nuclear cyclin B1 accumulation^9^, we also determined if the delayed cyclin B1 nuclear translocation of cells in hydrogels could be an indirect effect due to DNA damage, by measuring the levels of histone *γ*-H2AX. Neither confinement nor cell seeding on hydrogel increased the levels of *γ*-H2AX foci (Sup. Fig. 1e, f; ***p<0.001), suggesting that this process is independent of DNA damage.

**Figure 1.**
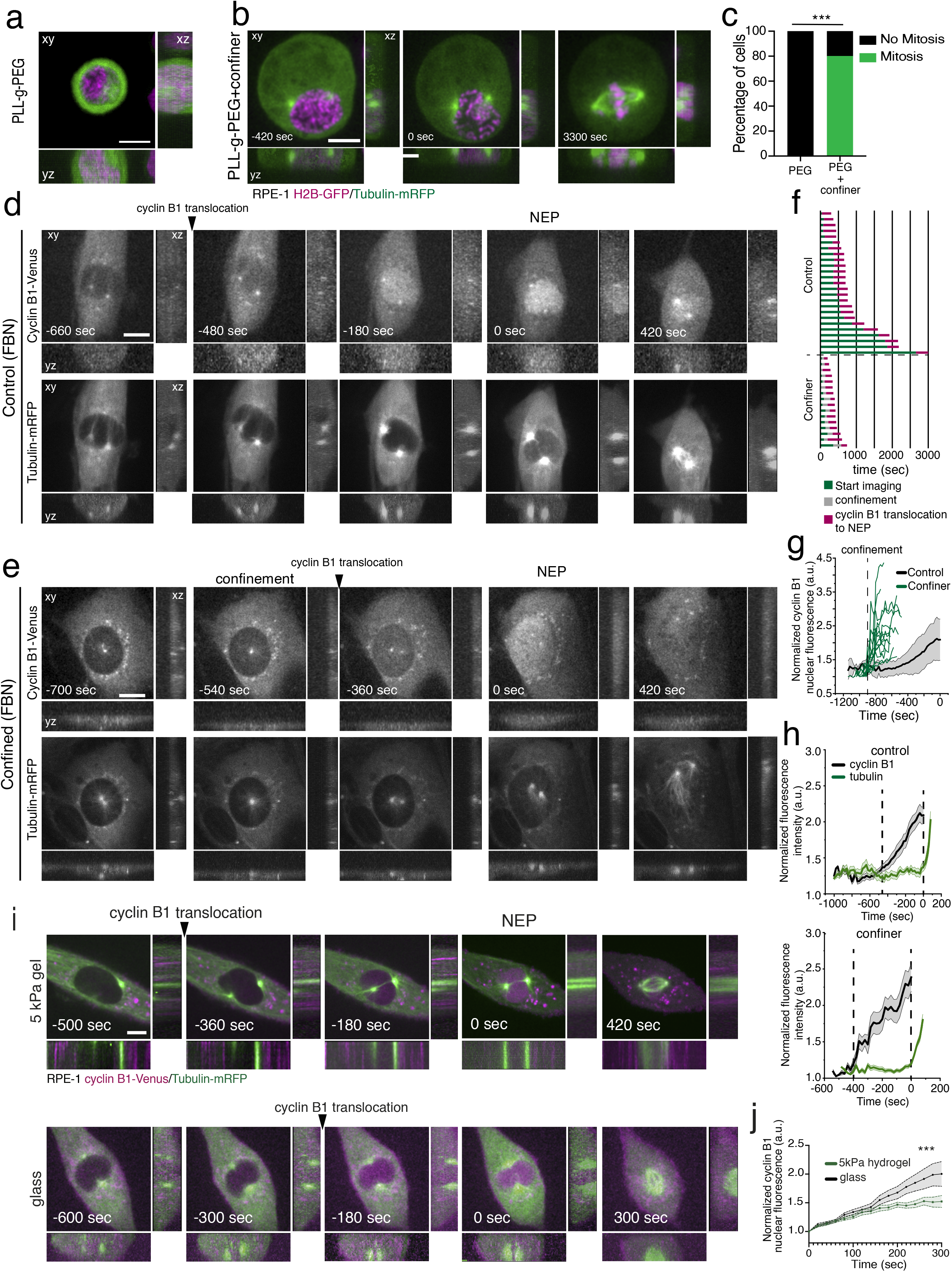
Cell confinement alters the nuclear translocation of cyclin B1. **(a)** Rpe-1 cell expressing H2B-GFP and tubulin-mRFP seeded on PLL-g-PEG coated substrate (non-adherent conditions). Scale bar corresponds to 10μm. **(b)** Rpe-1 cell expressing H2B-GFP and tubulin-mRFP dividing on a PLL-g-PEG coated substrate under geometric confinement. Time is in seconds and scale bar corresponds to 10μm. Images were acquired with a 20sec interval and time zero corresponds to NEP. **(c)** Percentage of cells that enter mitosis when seeded in non-adherent conditions with (n=10) and without (n=6; ***p<0.001) confinement. **(d)** Rpe-1 cell expressing cyclin B1-Venus (top panel) and tubulin-mRFP (lower panel) dividing on an FBN-coated substrate. Time is in seconds and scale bar corresponds to 10μm. Images were acquired with a 20sec interval and time zero corresponds to NEP. **(e)** Rpe-1 cell expressing cyclin B1-Venus (top panel) and tubulin-mRFP (lower panel) dividing on an FBN-coated substrate under confinement. Time frame in seconds and scale bar corresponds to 10μm. Images were acquired with 20sec interval and time zero corresponds to NEP. **(f)** Quantification of the time required for cyclin B1 nuclear accumulation and tubulin nuclear translocation in control (n=26) versus confined (n=19) conditions. **(g)** Normalized cyclin B1 fluorescence accumulation, inside the nucleus, over-time in control, non-confined cells (black; n=26) and individual tracks for confined cells (green; n=19; ***p<0.001). The vertical dashed line represents the moment of confinement. Time zero corresponds to NEP. **(h)** Correlation between the accumulation of cyclin B1 (black) and tubulin (green) in the nucleus over time in control cells (top panel) and confined cells (lower panel). Time zero corresponds to NEP. The vertical dashed lines represent the time interval between cyclin B1 nuclear entry and NEP. **(i)** Rpe-1 cell expressing cyclinB1-Venus and tubulin-mRFP dividing on a 5kPa hydrogel (top panel; n=16) or on glass (bottom panel; n=17). Time is in seconds and scale bar corresponds to 10μm. Images were acquired with 20sec interval and time zero corresponds to NEP. **(j)** Normalized cyclin B1 fluorescence accumulation inside the nucleus over-time in cells seeded on glass (black) or on a 5kPa hydrogel (green; ***p<0.001). Time zero corresponds to the lowest intensity value inside the cell nucleus.

This mechanical stimulation of cyclin B1 translocation was further confirmed by treating RPE-1 cells with a hypotonic shock (Sup. Fig. 2a, b), known to induce cell and nuclear membrane tension^28,29^, without generating apparent DNA damage^28^. As expected, the hypotonic shock induced a faster translocation of cyclin B1 into the nucleus, when compared to controls (Sup. Fig. 2a-c; ***p<0.001). Confinement of HeLa cells yielded similar faster translocation of cyclin B1 into the nucleus (Sup. Fig. 2d-f; **p<0.01), suggesting a conservation of this mechanism.

During mitotic entry, cells reorganize their cytoskeleton and round up^30^, decreasing the traction forces exerted by cells on the extracellular environment^26^. To test whether the process of cell rounding interferes with cyclin B1 translocation, we expressed a mutant form of the GTPase Rap1 (Rap1Q63E, hereafter named Rap1*), which effectively blocks focal adhesion disassembly and prevents cell rounding. Expression of Rap1* did not alter cyclin B1 translocation, even though cell rounding was efficiently blocked, as measured by cell membrane eccentricity^26^(Sup. Fig. 3a, b; ***p<0.001). Although we cannot completely rule out that cells with different rounding properties could exhibit changes in cyclin B1 translocation kinetics, our data is indicative that this process seems to be independent of cell rounding.

Next, we set to determine how this confinement-induced cyclin B1 translocation depended on the classical cyclin B1 import pathway. Because translocation in unconfined situations relies on cyclin B1-CDK1 activation^3^, we imaged cells treated with the CDK1 inhibitor RO-3306, with or without confinement (CDK1i; Fig. 2a-c). As expected, CDK1 inhibition effectively blocked cyclin B1 translocation to the nucleus (Fig. 2b, f). Interestingly, confinement was sufficient to overcome the inhibition of CDK1 and force translocation of cyclin B1 to the nucleus (Fig. 2c, f; ***p<0.001). However, these cells failed to enter mitosis, as NEP was blocked by CDK1 inhibition^31,32^. This observation further strengthens the idea that mechanical stimulation per se does not affect the nucleus barrier function. Accumulation of cyclin B1 relies on a balance between its nuclear import and export. Export is regulated by cyclin B1 binding to the exportin Crm1^33^, whereas import is dependent on binding to importin β^34^, and is greatly enhanced by phosphorylation of cyclin B1 on its CRS sequence^35,36^. We started by treating cells with importazole, to inhibit importin β function. This treatment efficiently blocked cyclin B1 nuclear translocation, even in confinement conditions (Fig. 2d, g), indicating that the accelerated translocation of cyclin B1 to the nucleus cannot be explained by an increased diffusive shuttling alone, but by an active process, dependent on importin β. A similar block in cyclin B1 translocation was observed when cells expressing a mutant form of cyclin B1 with its five CRS phosphorylation sites mutated to alanines (cyclin B1-5A-GFP)^35^, were subjected to confinement (Sup. Fig. 3c, d). On the other hand, when nuclear export was inhibited by treatment with Leptomycin B we observed a delay in nuclear accumulation of cyclin B1 (Fig. 2e, h), in accordance with previous reports^37^, that was only partially rescued by confinement (Fig. 2h; ***p<0.001). In addition to the mechanisms described above, Plk1 activity has also been recently proposed to regulate mitotic entry and cyclin B1-CDK1 activity^38^. To determine if confinement-induced translocation was dependent on Plk1 activity, we treated RPE-1 cells with 200nM of the Plk1 inhibitor BI2536 for 2h (Plk1i), a dose previously reported to block mitotic entry^38^. Accordingly, this treatment prevented cells from accumulating cyclin B1 in the nucleus and entering mitosis (Sup. Fig. 3e, g, h). Strikingly, confining Plk1 i cells was sufficient to rescue the accumulation of cyclin B1 and allow mitotic entry (Sup. Fig. 3f-h; ***p<0.001). It should be noted that confined, Plk1i-treated cells that failed to enter mitosis also did not accumulate cyclin B1 in the nucleus (Sup. Fig. 3g, h). Overall, our data strongly suggest that a mechanical signal acts in parallel with the biochemical pathways to help regulate the timing of cyclin B1 nuclear accumulation and control mitotic entry, in a process that requires phosphorylation of cyclin B1 in the CRS and binding to importin β.

**Figure 2.**
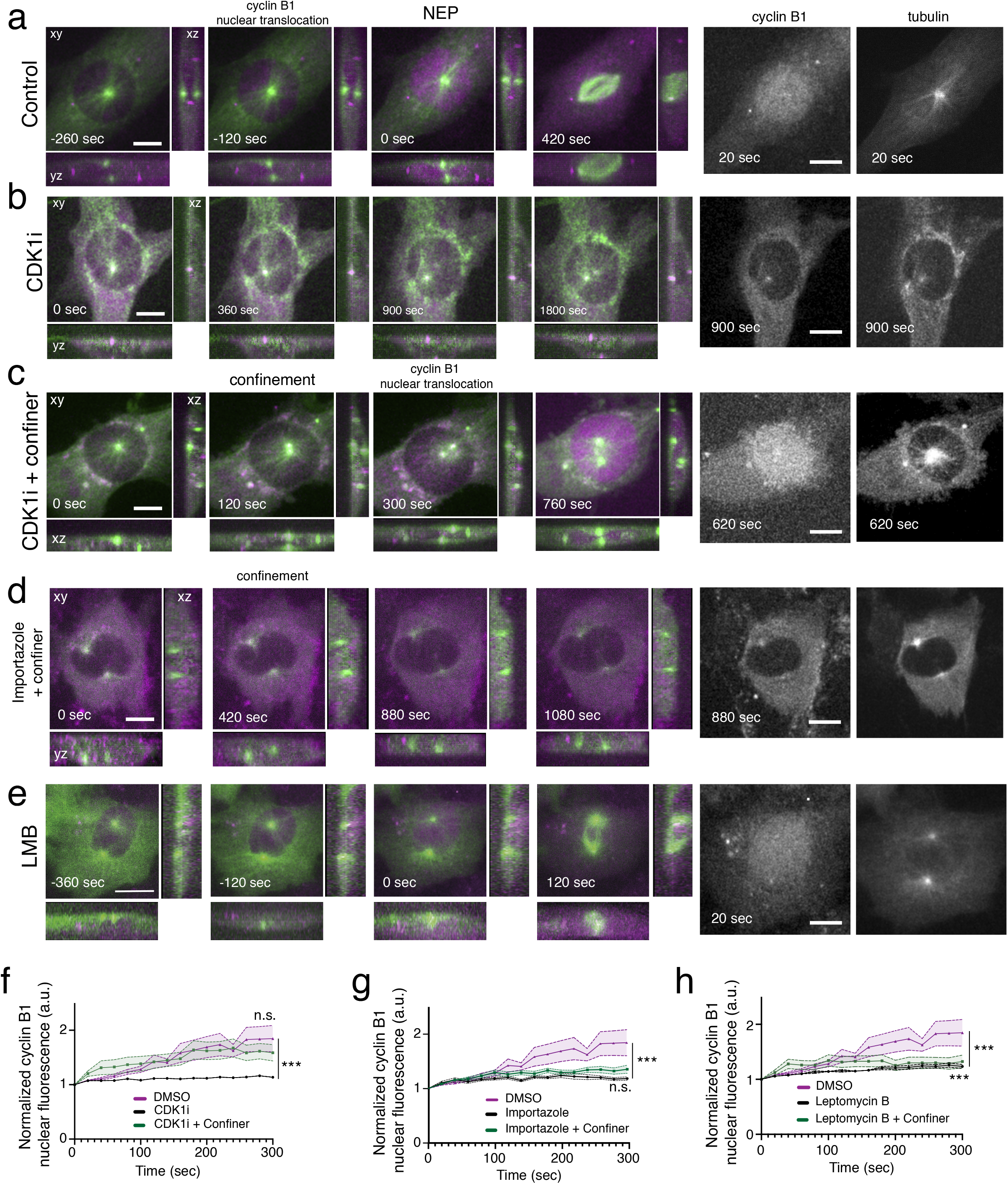
Characterization of confinement-induced cyclin B1 translocation. **(a)** RPE-1 cell expressing cyclinB1-Venus and tubulin-mRFP, dividing on a FBN-coated substrate. Right panels highlight cyclin B1 and tubulin accumulation at the nucleus. Time frame in seconds and scale bar corresponds to 10μm. Images were acquired with 20sec interval and time zero corresponds to NEP. **(b)** Rpe-1 cell expressing cyclinB1-Venus and tubulin-mRFP seeded on an FBN-coated substrate, treated with a CDK1 inhibitor (CDK1i). Right panels highlight the lack of cyclin B1 and tubulin accumulation inside the nucleus. Time frame in seconds and scale bar corresponds to 10μm. Images were acquired with 20sec interval and time zero corresponds to the first frame. **(c)** RPE-1 cell expressing cyclinB1-Venus and tubulin-mRFP seeded on a FBN-coated substrate, treated with CDK1i and under confinement. Right panel shows confinement-induced nuclear accumulation of cyclin B1 and the lack of nuclear tubulin accumulation. Time frame in seconds and scale bar corresponds to 10μm. Images were acquired with 20sec interval and time zero corresponds to the first frame. **(d)** RPE-1 cell expressing cyclinB1-Venus and tubulin-mRFP seeded on a FBN-coated substrate, treated with importazole and under confinement. Right panel highlights the absence of nuclear accumulation of cyclin B1 and tubulin. Time frame in seconds and scale bar corresponds to 10μm. Images were acquired with 20sec interval and time zero corresponds to the first frame. **(e)** RPE-1 cell expressing cyclin B1-Venus and tubulin-mRFP seeded on an FBN-coated substrate, treated with Leptomycin B1 (LMB; n=19). Time frame in seconds and scale bar corresponds to 10μm. Images were acquired with 2Osec interval and time zero corresponds to NEP. Right panels highlights the nuclear accumulation of cyclin B1 and tubulin. **(f)** Normalized nuclear cyclin B1 fluorescence over time in control, DMSO-treated cells (magenta; n=17), cells treated with CDK1i (black; n=23) and cells treated with CDK1i and subjected to confinement (green; n=17; ***p<0.001). Time zero corresponds to the lowest fluorescence intensity value inside the cell nucleus. **(g)** Normalized nuclear cyclin B1 fluorescence over time, in cells treated with importazole (black; n=27), cells treated with importazole and under confinement (green; n=17) and control, DMSO-treated cells (magenta; n=17). Time zero corresponds to the lowest fluorescence intensity value inside the cell nucleus. (***p<0.001; n.s. – not significant). **(h)** Normalized nuclear cyclin B1 fluorescence over time for cells treated with Leptomycin B1 (black; n=19), cells treated with Leptomycin B1 under confinement (green; n=16) and control DMSO treated cells (magenta; n=17; p<0.001). Time zero corresponds to the lowest intensity value inside the cell nucleus. Note the DMSO-treated group is the same for panels f, g and h. Statistical analysis of cyclin B1 accumulation was performed for the entire time course using an ANOVA Repeated Measures test when samples had a normal distribution. Otherwise, analysis was done using a Repeated Measures ANOVA on Ranks.

We then sought to identify potential mechanosensing mechanisms involved in the transport of cyclin B1 to the nucleus. Mechanical forces generated by the cytoskeleton are transmitted to the nucleus through the linker of cytoskeleton and nucleoskeleton (LINC) complex^18^. To address how force transmission might affect cyclin B1 nuclear translocation, we expressed a dominant-negative form of nesprin (DN-KASH) that prevents its binding to SUN proteins and blocks force propagation^18^. Expression of DN-KASH significantly delayed cyclin B1 nuclear accumulation (Fig. 3a, i; ***p<0.001). Similarly, we observed delays in cyclin B1 translocation following inhibition of ROCK with Y-27632 (Fig. 3c, j; **p<0.01), depletion of ROCK1 with shRNA (Fig. 3f, k; ***p<0.001), myosin II inhibition with para-nitro-blebbistatin (p-N-blebb; Fig. 3e, l; **p<0.01), inhibition of myosin light chain kinase (MLCK) with ML-7 (Fig. 3k), and actin depolymerization with cytochalasin D (CytoD; Fig. 3g, m; ***p<0.001). Conversely, microtubule depolymerization with nocodazole did not affect cyclin B1 translocation (Noc; Fig. 3h, n; n.s. – not significant). Importantly, confinement was able to fully rescue cyclin B1 accumulation that was lost upon expression of DN-KASH (Fig. 3b, i; ***p<0.001) or treatment with Y27632 (Fig. 3d, h), and partially rescued accumulation following actin (Fig. 3f; ***p<0.001) or myosin (Fig. 3g; **p<0.01) inhibitions. Overall, these experiments identify actomyosin-dependent force transmission to the nucleus as important players in facilitating cyclin B1 nuclear translocation.

**Figure 3.**
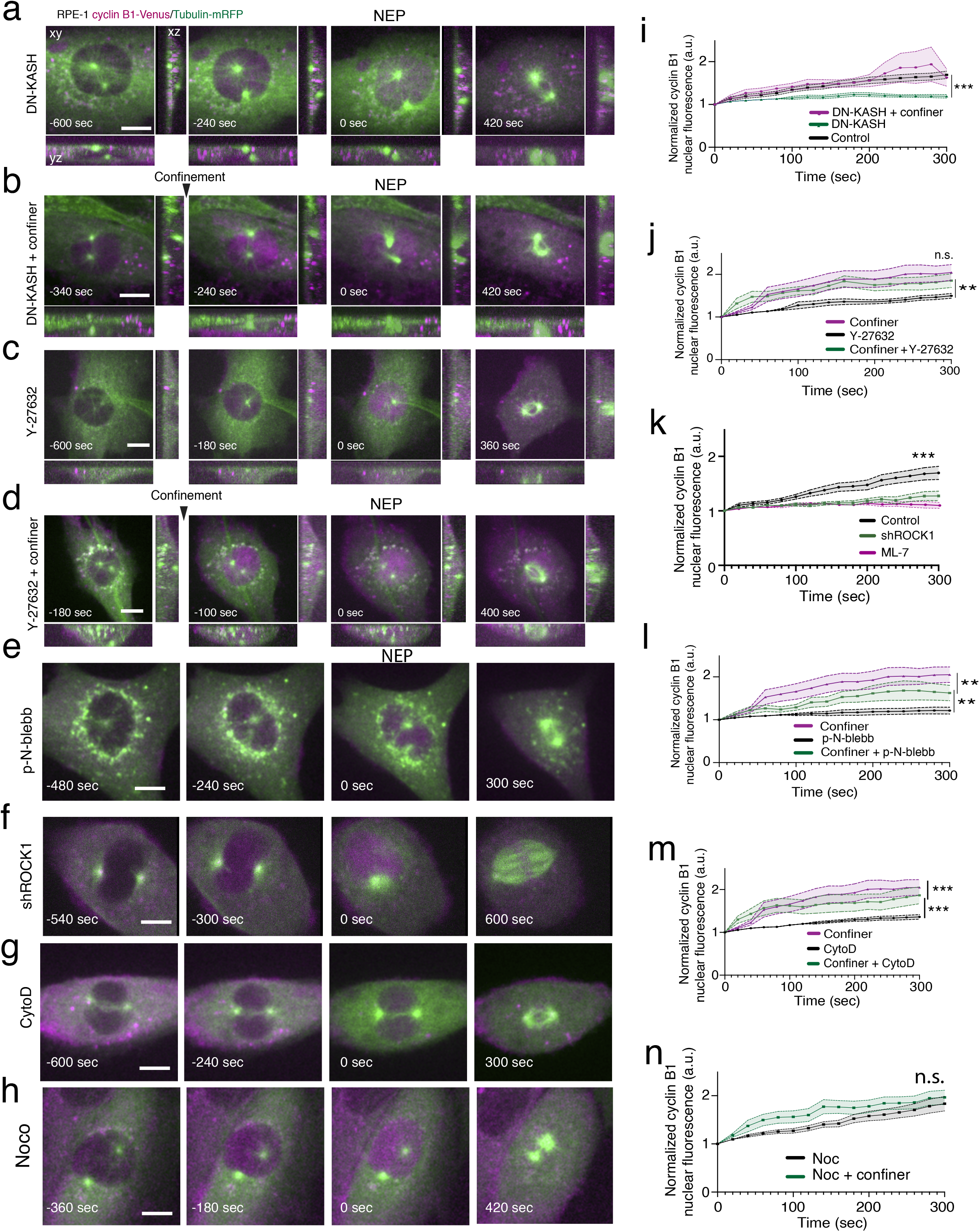
Actomyosin contractility contributes to cyclin B1 translocation. **(a)** RPE-1 cell expressing cyclinB1-Venus and tubulin-mRFP dividing on an FBN-coated substrate, expressing a DN-KASH mutant. **(b)** RPE-1 cell expressing cyclin B1-Venus and tubulin-mRFP dividing on an FBN-coated substrate, expressing a DN-KASH mutant and subjected to confinement. **(c)** RPE-1 cell expressing cyclin B1-Venus and tubulin-mRFP dividing on an FBN-coated substrate and treated with Y-27632. **(d)** RPE-1 cell expressing cyclin B1-Venus and tubulin-mRFP dividing on an FBN-coated substrate, treated with Y-27632 and dividing under confinement. **(e)** RPE-1 cell expressing cyclin B1-Venus and tubulin-mRFP dividing on an FBN-coated substrate and treated p-Nitro-blebbistatin (p-N-blebb; n=15). **(f)** RPE-1 cell expressing cyclin B1-Venus and tubulin-mRFP dividing on an FBN-coated substrate and depleted for ROCK1 (shROCK1). **(g)** RPE-1 cell expressing cyclin B1-Venus and tubulin-mRFP dividing on an FBN-coated substrate and treated Cytochalasin D (CytoD). **(h)** RPE-1 cell expressing cyclin B1-Venus and tubulin-mRFP dividing on an FBN-coated substrate and treated Nocodazole (Noco). For all images, time frame in seconds and scale bar corresponds to 10μm. Images were acquired with a 20sec interval and time zero corresponds to NEP. **(i)** Normalized nuclear cyclinB1 fluorescence accumulation over time for non-confined cells expressing the DN-KASH mutant (green; n=17), confined cells expressing the DN-KASH mutant (magenta; n=17) and control cells treated with Lipofectamin 2000 (black; n=22). Time zero corresponds to the lowest intensity value inside the cell nucleus (***p<0.001). **(j)** Normalized nuclear cyclin B1 fluorescence accumulation over time in non-confined cells treated with Y-27632 (black; n=21), confined cells treated with Y-27632 (green; n=19) and confined cell only (magenta; n=14). Time zero corresponds to the lowest intensity value inside the cell nucleus (**p<0.01). **(k)** Nuclear accumulation of cyclin B1 fluorescence accumulation over time, for control cells (black; ***p<0.001), cells expressing shROCK1 (green) and cells treated with ML-7 (magenta). Time zero corresponds to the lowest value of fluorescence inside the nucleus. **(l)** Normalized nuclear cyclinB1 fluorescence accumulation over time in non-confined cells treated with p-Nitro-blebbistatin (p-N-blebb; black; n=15), confined cells treated with p-N-blebb (green; n18) and confined cells (magenta; n=14). Time zero corresponds to the lowest intensity value inside the cell nucleus (**p<0.01). **(m)** Normalized nuclear cyclin B1 fluorescence accumulation over-time in non-confined cells treated with Cytochalasin D (CytoD; black; n=26), confined cells treated with CytoD (green; n=18) and confined cells only (magenta; n=14). Time zero corresponds to the lowest intensity value inside the cell nucleus (***p<0.001). **(n)** Normalized nuclear cyclin B1 fluorescence accumulation over time, for cells treated with Noco without (black n= 20) or with confinement (green n=20; n.s. – not significant). Note the confiner only group is the same for panels j, l and m. Statistical analysis of cyclin B1 accumulation was performed for the entire time course using an ANOVA Repeated Measures test when samples had a normal distribution. Otherwise, analysis was done using a Repeated Measures ANOVA on Ranks.

Our results indicate that during the G2-M transition, the NE unfolds (Sup. Fig. 1), an event that can be exacerbated by mechanical confinement (Sup. Fig. 1). Moreover, we showed that actomyosin contractility facilitates the translocation of cyclin B1 into the nucleus (Fig. 3). Since increased actomyosin contractility was recently shown to induce NE unfolding^21,25^, we tested whether such a mechanism also acted during the G2-M transition. To do so, we evaluated the nuclear irregularity index (NII) of interphase and prophase nuclei using fixed-cell analysis. This parameter was calculated as 1-solidity (solidity is defined as nucleus area/nucleus convex area). Our results confirm a decrease in NII in prophase cells, when compared to interphase, indicating an unfolding of the NE (Fig. 4a-c; ***p<0.001). Nuclear unfolding was previously associated with increased nuclear tension^39^, which triggers the recruitment and activation of the calcium-dependent, nucleoplasmic phospholipase cPLA2 to the NE. Active cPLA2 then stimulates actomyosin contractility^21,25^, creating a positive feedback loop. If indeed prophase nuclei are under increased tension, it is possible that cPLA2 is recruited to the NE at this stage. Accordingly, we found that cPLA2 is recruited to the NE during prophase, similarly to what happens in confined interphase cells^21,25^ (white arrows; Fig. 4d, e; ***p<0.001), suggestive of increased NE tension and cPLA2 activation at this stage. While our data shows that NE unfolding during prophase can recruit cPLA2, it does not explain how the NE unfolds in the first place. To determine this, we analysed the NII of cells expressing DN-KASH or treated with actomyosin inhibitors. All treatments led to an increase in NII, which was reverted upon confinement (Fig. 4f, g and Sup. Fig. 4a; ***p<0.001). This confinement-generated decrease in NII likely reflects an unfolding of the NE, which is evident from the images of confined nuclei (Fig. 4f and Sup. Fig. 4a) as well as the increased NPC-NPC distance (Sup. Fig. 4b, c; ***p<0.001). Collectively, these data indicate that actomyosin contractility triggers NE unfolding during prophase. Importantly, blocking actomyosin or expressing DN-KASH significantly decreased cPLA2 accumulation on the NE (Fig. 4f, h and Sup. Fig. 4a; ***p<0.001), which was not rescued by confinement (Fig. 4h; ***p<0.001). This indicates cPLA2 recruitment to the NE requires an intact connection between the cytoskeleton and nucleus. Overall, we conclude that an increase in actomyosin contractility during prophase is required to unfold the NE, increasing nuclear tension, and leading to cPLA2 recruitment. The question remains of the functional relevance of cPLA2 NE recruitment. If cPLA2 is functionally important to facilitate cyclin B1 translocation, inhibiting its activity should result in a delay in cyclin B1 nuclear accumulation. Indeed, inhibition of cPLA2 activity with AAOCF3 led to a significant decrease in cyclin B1 nucleoplasmic shuttling (Fig. 4i and Sup. Fig. 4d, e; **p<0.01; ***p<0.001). Similarly, acutely interfering with the release of internal Ca^2+^ stores (known to trigger cPLA2 NE recruitment and activation^39^), using BAPTA-AM + 2APB also decreased cyclin B1 nuclear translocation (Fig.4j; Sup Fig. 4d, e; ***p<0.001), as anticipated. While we cannot rule out that interfering with calcium release might affect other cellular processes, the use of BAPTA to disrupt internal calcium release during prophase has been previously described^40^ and should not affect NEP at the concentrations used in our study. Moreover, we sought to minimize possible side-effects by adding the drugs acutely in late G2. Remarkably, confinement was able to stimulate cyclin B1 translocation, even when cPLA2 activity or calcium release were inhibited (Fig. 4i, j; **p<0.01; ***p<0.001). This likely occurs due to confinement-induced unfolding of the NE (Sup. Fig. 1), that is sufficient to bypass the pharmacological inhibition of contractility and still induce an increase in NPC distance (Sup. Fig. 4b, c; ***p<0.001). Together, these observations support a working model for the mechanical regulation of mitotic entry based on actomyosin activity, that triggers nuclear unfolding, leading to cPLA2 activation. This facilitates cyclin B1 transport across the NPCs, increasing its nuclear accumulation.

**Figure 4.**
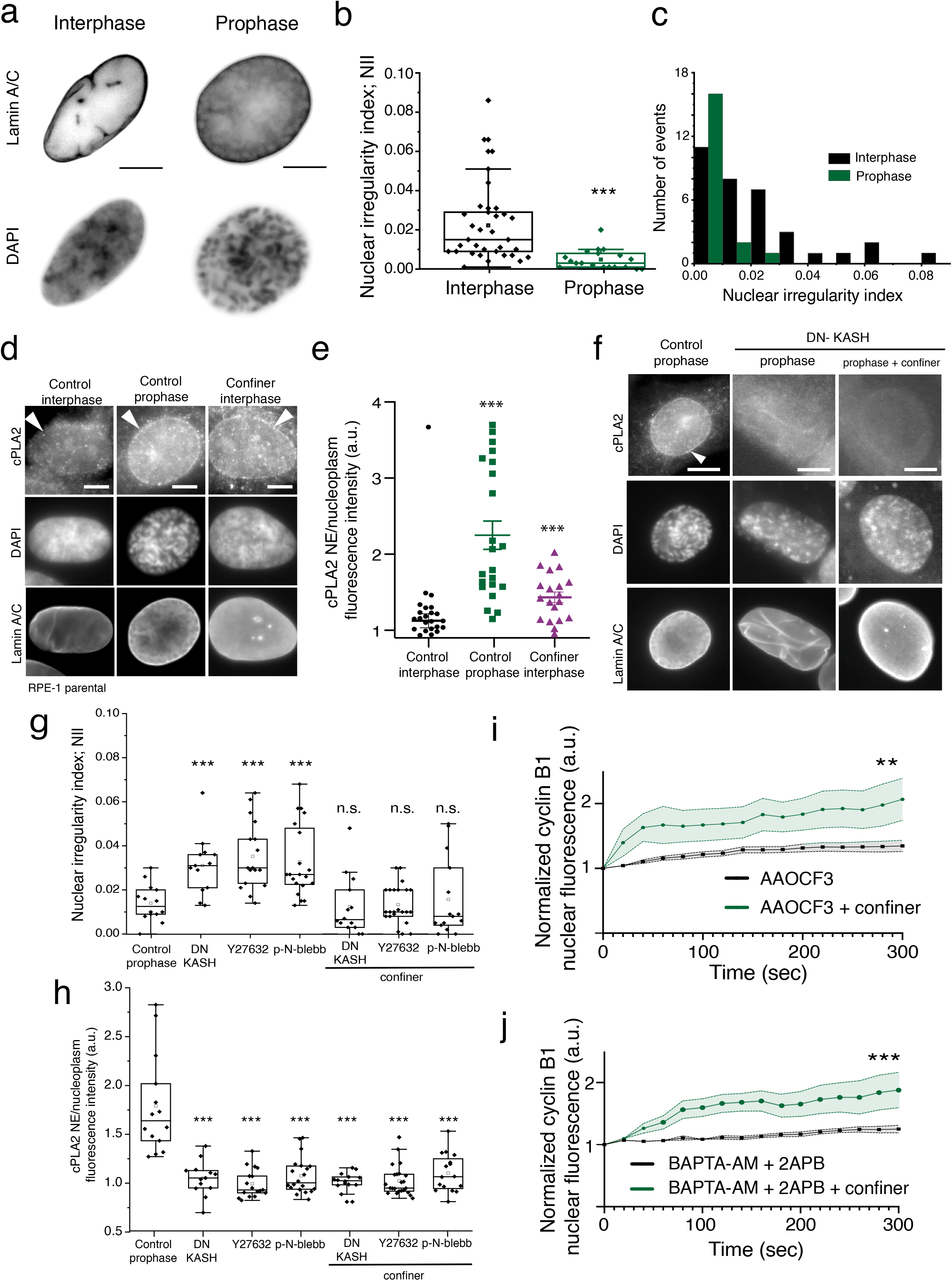
Nuclear unfolding during prophase recruits cPLA2 to the NE. **(a)** Representative images of the nucleus of interphase and prophase cells immunostained for Lamin A/C and DAPI. Please note the irregularities on the NE surface in interphase cells. Scale bars, 10μm. **(b)** Nuclear irregularity index (NII) in interphase (black; n=34) versus mitotic cells (green; n=20; ***p<0.001). **(c)** Distribution of NII values in interphase (black) and mitotic cells (green). **(d)** Representative immunofluorescence images of parental RPE-1, interphase non-confined cells (control interphase; n=32), non-confined prophase cells (control prophase; n=28) and interphase confined cells (interphase confiner; n=19) stained for cPLA2, Lamin A/C and DAPI. Scale bars, 10μm. **(e)** cPLA2 fluorescence intensity ratio between the NE and the nucleoplasm for control interphase cells (black), control prophase cells (green; ***p<0.001) and confiner interphase cells (magenta; ***p<0.001). Comparison between experimental groups was performed using a One-way ANOVA test. **(f)** Representative immunofluorescence images of parental Rpe-1 cells in prophase, non-confined conditions (Control prophase; n=14), prophase cells expressing the DN-KASH (n=26) and prophase cells expressing DN-KASH under confinement (n=27), stained with cPLA2, DAPI and Lamin A/C. **(g)** Nuclear irregularity index (NII) in control prophase cells (n=15), prophase cells expressing DN-KASH without (n=14) or with (n=13) confinement, prophase cells treated with Y-27632 without (n=17 or without (n=23) confinement or cells treated p-Nitro-blebb without (n=20) and with confinement (n=16; ***p<0.001; n.s. – not significant). **(h)** cPLA2 fluorescence intensity ratio between the NE and the nucleoplasm for control prophase cells, prophase cells expressing DN-KASH, treated with Y-27632 or treated with p-Nitro-blebb without and with confinement (***p<0.001). **(i)** Normalized nuclear cyclinB1 fluorescence accumulation over time in non-confined cells treated with cPLA2 inhibitor (AACOCF3, black; n=15) and confined cells treated with cPLA2 inhibitor (green; n=12). Time zero corresponds to the lowest fluorescence intensity value inside the nucleus (**p<0.01). **(j)** Normalized nuclear cyclinB1 fluorescence accumulation over time in non-confined cells treated with BAPTA-AM + 2APB (black; n=17) and confined cells treated with BAPTA-AM + 2APB (green; n=11). Time zero corresponds to the lowest fluorescence intensity value inside the nucleus (***p<0.001). Statistical analysis of cyclin B1 accumulation was performed for the entire time course using an ANOVA Repeated Measures test when samples had a normal distribution. Otherwise, analysis was done using a Repeated Measures ANOVA on Ranks.

Nuclear translocation of cyclin B1 sets the time for the G2-M transition^41^ and is essential for preventing untimely mitotic entry, which results in chromosome segregation errors^42^. Similarly, confining cells throughout mitosis also contributes to the occurrence of segregation errors^1,43^. Whether a short confinement during prophase, which is sufficient to induce premature cyclin B1 translocation and NEP, results in chromosome segregation errors remains unknown. To test this, we subjected cells in prophase to a short confinement, which was released shortly after NEP (Fig. 5a). This approach should induce mitotic entry, while still providing enough volume for the spindle to assemble unconstrained^1^. Cells were then allowed to progress through mitosis unperturbed, so that we could determine mitotic timings, as well as the rate of chromosome missegregation. Notably, a significant proportion of cells that were subjected to short confinement entered mitosis with incomplete centrosome separation (Fig. 5a-d), which we and others have shown^26,44^ can increase the frequency of mitotic errors. Detailed analysis of centrosome behaviour during the early stages of spindle assembly under short confinement also revealed that centrosome separation and positioning still depended on kinesin-5 and dynein activities. Accordingly, mechanical confinement could not rescue the monopolar spindles generated by STLC treatment (Sup. Fig. 5a, b) or the centrosome positioning defects induced by DHC RNAi (Sup. Fig. 5c-e). As a result, this acute confinement resulted in increased chromosome segregation errors (Fig. 5e; *p<0.05) and a slight mitotic delay (Fig. 5f; 24±7 min for controls vs. 36±20 min for confined cells; *p<0.05), when compared to unconfined cells. It is plausible that these errors arise from a confinement-induced acceleration of NEP, preventing cells from properly organizing a mitotic spindle. To test this, we decided to generate an artificial rupture of the NE with laser microsurgery^45^, allowing cyclin B1 and tubulin to enter the nuclear space and anticipating mitotic entry (Fig. 5g). Using this approach, we triggered immediate mitotic entry, which was sufficient to increase chromosome missegregation events (white arrowhead, Fig. 5g, h; **p=0.02) and induce a slight mitotic delay (Fig. 5i). Together, these experiments demonstrate that untimely mitotic entry through acute mechanical confinement during the G2-M transition, can have deleterious downstream consequences for chromosome segregation.

**Figure 5.**
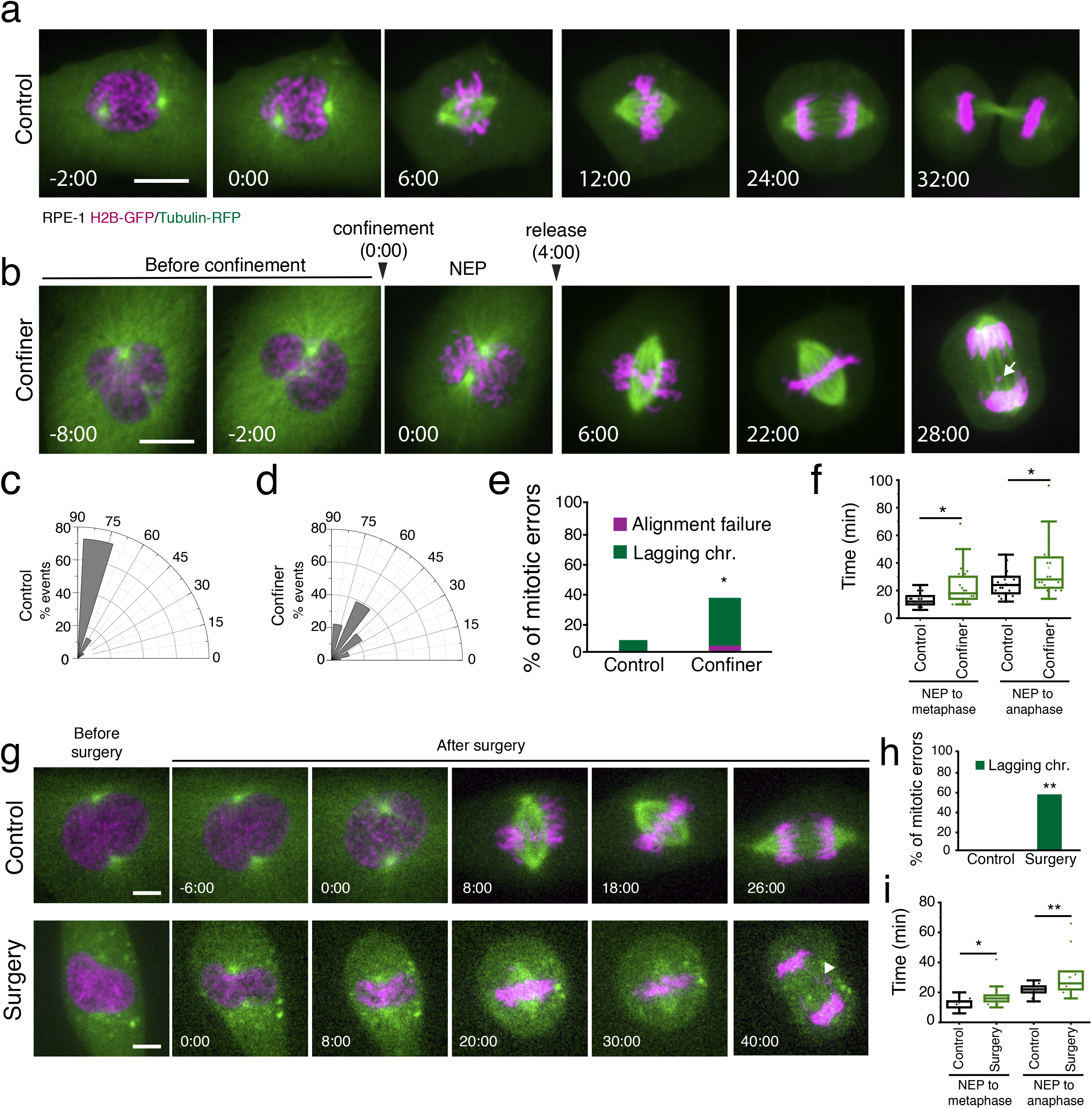
Premature mitotic entry potentiates chromosome segregation errors. RPE-1 cell expressing H2B-GFP and tubulin-mRFP dividing on an FBN-coated substrate without **(a)** or with short-term confinement **(b)**. Time frame in min:sec and scale bars corresponds to 10μm. Images were acquired with a 2min interval and time zero corresponds to NEP. Inter-centrosomes angle at the moment of NEP in cells dividing without **(c)** or with confinement **(d). (e)** Percentage of mitotic errors in control, non-confined cells (n=26) and cells under a short period of confinement (n=23; *p<0.05). **(f)** Mitotic timings (NEP-metaphase and NEP-anaphase) in control, non-confined cells (black) and cells under a short period of confinement (green; *p<0.05). **(g)** RPE-1 cell expressing H2B-GFP and tubulin-mRFP dividing on an FBN-coated substrate in control conditions, where laser microsurgery was performed on the cytoplasm (top panel; n=12), conditions where the NE was ruptured using laser microsurgery (lower panel; n=15). Time frame in seconds and scale bar corresponds to 10μm. Images were acquired with 2min interval and time zero corresponds to NEP. **(h)** Percentage of mitotic errors in control cells and cells where the NE was ruptured using laser microsurgery (**p<0.01). **(i)** Mitotic timings (NEP-metaphase and NEP-anaphase) in control cells (black) and cells where the NE was ruptured using laser microsurgery (green; *p<0.05; **p<0.01).

The biochemical regulation of the G2-M transition has been studied extensively^3,8–10,35,36^. However, how mechanical forces affect this essential step of the cell cycle was unknown. Here, we propose a nongenetic, mechanical pathway based on nuclear tension that acts during the G2-M transition, impacting cyclin B1 translocation and NEP. In agreement with this model, we observed an actomyosin-dependent nuclear unfolding in prophase cells (Fig. 4a-c; Supplementary Figs. 1 and 4), similarly to previous observations in G2 cells^21^. This unfolding increases cPLA2 recruitment to the NE (Fig. 4d, e), indicative of cPLA2 activation^21,25,39^ and likely reflect an increased tension on the nucleus during the G2-M transition. It is tempting to speculate that during the G2-M transition, this increased nuclear tension could be sufficient to deform the nucleus and NPCs, leading to faster cyclin B1 transport across the NE, as was proposed for YAP or MyoD ^20,22^. In fact, such tension was recently shown to deform the NPCs in vivo, and facilitate transport across the NE^46^. In agreement, we were able to rescue cyclin B1 shuttling to the nucleus by confinement, even in the absence of cPLA2 activity or actomyosin contractility. Overall, we propose this mechanical pathway cooperates with the classical cyclin B1 transport machinery to fine tune NEP according to the cellular tension state, thus ensuring timely and accurate cell division. Accordingly, confinement could not induce cyclin B1 translocation in the absence of importin β activity (Fig. 2d, f) or CRS phosphorylation (Sup. Fig. 3c, d).

Establishing mechanical forces as an important player in cyclin B1 nuclear translocation and mitotic entry raises the interesting possibility that the nucleus might act as a sensor^24,28^ for external forces, regulating cell cycle progression and cell division to control tissue growth and avoid over-proliferation.

## Materials and Methods

### Cell lines

Cell lines used were cultured in DMEM (Life Technologies) supplemented with 10% FBS (Fetal Bovine Serum – Life Technologies) and kept in culture in a 37°C humidified incubator with 5% CO_2_. RPE-1 parental and RPE-1 H2B-GFP and tubulin-mRFP were already available in our lab. RPE-1 endogenous cyclinB1-Venus cell line was a gift from Jonathon Pines. RPE-1 cyclinB1-Venus/tubulin-mRFP cell line was generated in our lab by transduction with lentiviral vectors containing tubulin-mRFP (Addgene). RPE-1 TUB-mRFP cells were generated in our lab by transduction with lentiviral vectors containing tubulin m-RFP (Addgene). HEK293T cells at a 50-70% confluence were co-transfected with lentiviral packaging vectors (16.6 μg of Pax2, 5.6 μg of pMD2 and 22.3 μg of LV-tubulin-mRFP), using 30μg of Lipofectamin 2000 (Life Technologies). Approximately 4-5 days after the transduction, the virus-containing supernatant was collected, filtered, and stored at −80°C. RPE-1 cyclinB1-Venus cells were infected with virus particles together with polybrene (1:1000) in standard culture media for 24 h. Approximately 2-3 days after the infection, the cells expressing tubulin were isolated by fluorescence activated cell sorting (FACS; FACS Aria II). To deplete ROCK1, a shRNAi-LV was co-transfected with a lentiviral packaging vector as previously explained. The viral particles were then transduced in the RPE-1 Tub-mRFP cell line to generate stable cells depleted of ROCK1.

### Drug treatments

CDK1 inhibitor (RO-3306) was used at a concentration of 9 μM for 16 h. Importazole was added to the cells at a final concentration of 40 μM 2 h before the experiment, ROCK inhibitor (Y-27632) was used at 5 μM for 30 min (Sigma Aldrich). To interfere with cPLA2 activity, AACOCF3 was used at 20 μM (TOCRIS) for 30 min. To block the release of calcium ions from internal cellular stores, BAPTA-AM and 2APB (Abcam) were used at 10 μM for 15-30 min. Myosin activity was perturbed using p-nitro-blebbistatin at 50 μM for 30 min (MotorPharma). MLCK activity was blocked using ML-7 at 50 μM for 30 min. To interfere with the actin cytoskeleton, we used cytochalasin D at 0.5 μM (TOCRIS) for 30 min. To perturb microtubules, nocodazole was used at 3.3 μM for 30 min. Plk1 inhibitor (BI2536), from Axon Med Chem, was used at 200mM for 2h. To induce DNA damage, cells were treated with 1μM of Etoposide (Selleck Chemicals CO., Ltd) for 2h before fixation. Control cells were treated with either DMSO (Sigma) or transfected with Lipofectamin 2000 (Invitrogen), as explained in the text.

### Hypotonic Shock

To perform the hypotonic shock, RPE-1 cells expressing cyclinB1-Venus and tubulin-mRFP were seeded as described above. At the microscope, MiliQ water was added to the imaging medium (1:5 dilution).

### Transfections

Cells were transfected with the plasmid encoding the DN-KASH or the Rap1Q63E (Rap1*) mutant using Lipofectamin 2000 (Life Technologies).

Specifically, 5 μL of Lipofectamin 2000 and 0.5 μg of plasmid DNA were diluted separated and incubated in OPTIMEM (Alfagene) for 30 min. The mixture was then added to confluent cells cultured and incubated for 6 h in reduced serum medium (DMEM with 5%FBS). Cells were analyzed 48 h after transfection.

For the DHC RNAi experiment, cells were transfected with small interfering RNAs (siRNAs) using Lipofectamin RNAi Max (Life Technologies). Specifically, 5 μL of Lipofectamin and 20 nM of each siRNA were diluted separated and incubated in OPTIMEM (Alfagene) for 30 min. The mixture was then added to confluent cells cultured and incubated for 6h in reduced serum medium (DMEM with 5% FBS). Commercial ON-TARGETplus SMARTpool siRNAis (Dharmacon) were used. Cells were analyzed 72 h after transfection and protein depletion efficiency verified by immunoblotting.

### Time-lapse microscopy

Between 12-24 h before the experiments 1.5×10^5^ cells were seeded on fluorodishes (WPI) coated with FBN (25 μg/mL; F1141; Sigma). Shortly before each experiment, DMEM 10%FBS medium was changed to Leibovitz’s-L15 medium (Life Technologies), supplemented with 10% FBS and Antibiotic-Antimycotic 100X (AAS; Life Technologies). Live cell imaging experiments were performed using temperature-controlled Nikon TE2000 microscopes equipped with a modified Yokogawa CSU-X1 spinning-disk head (Yokogawa Electric), an electro multiplying iXon+ DU-897 EM-CCD camera (Andor) and a filter wheel. Three laser lines were used to excite 488 nm, 561 nm and 647 nm and all the experiments were done with oil immersion 60x 1.4NA Plan-Apo DIC (Nikon). Image acquisition was controlled by NIS Elements AR software. Images with 17-21 z-stacks (0.5 μm step) were collected with a 20 sec interval.

### Western Blotting

Cell extracts were collected after trypsinization and centrifuged at 1,200 rpm for 5min, washed and resuspended in 30-50 μL of lysis buffer (20nM HEPES/KOH, pH 7.9; 1mM EDTA pH8; 150mM NaCl; 0.5% NP40; 10% glycerol, 1:50 protease inhibitor; 1:100 Phenylmethylsulfonul fluoride). The cells were then flash frozen in liquid nitrogen and kept on ice for 30 min. After centrifugation at 14,000 rpm for 8min at 4°C, the supernatant was collected, and protein concentration determined using the Bradford protein assay (Bio-Rad). The proteins were run on a 10% SDS-PAGE gel (50 μg/lane) and transferred using a wet blot apparatus for 1.5h at 70 V, with constant amperage. Later, the membranes were blocked with 5% milk in Tris Buffered Saline (TBS) with 0.1% Tween-20 (TBS-T) for 1h at room temperature. The primary antibodies used were anti-dynein heavy chain (1:250, Thermo Fisher Scientific) and anti-vinculin (1:1000, Thermo Fisher Scientific). All primary antibodies were incubated overnight at 4°C with shaking. After three washes in TBS-T, the membranes were incubated with the secondary antibody for 1 h at room temperature. The secondary antibodies used were anti-rabbit-HRP at 1:5000. After several washes with TBS-T, the detection was performed with Clarity Western ECL Substrate (Bio-Rad).

### Cell confinement setup

For dynamic confinement experiments, we adapted a cell confiner as previously described^27^, using a custom-designed polydimethylsiloxane (PDMS, RTV615, GE) layout to fit a 35 mm fluorodish. A suction cup was custom-made with a 10/1 mixture (w/w PDMS A/crosslinker B), baked on an 80°C hot plate for 1h and left to dry over-night before unmoulding. The confinement slide was polymerized on 10mm round coverslips. These round coverslips were first treated with air plasma for 2 min (Zepto system, Diener Electronics) and incubated with a 0.3% Bind-Silane (Sigma M6514)/5% acetic acid solution in ethanol. Then, the coverslips were rinsed with ethanol and left to dry. A gel with approximately 15kPa stiffness was prepared using an acrylamide (Bio-Rad)/bisacrylamide (Bio-Rad) mix. The mixture was added to the coverslips and allowed to polymerize. After polymerization, gels were hydrated with PBS and incubated with cell culture medium for at least 30 min. The confinement slide was then attached to the PDMS suction cup described above and connected to a vacuum generator apparatus (Elveflow).

For static confinement experiments, we used a commercially available 6-well confinement device (4DCell) with custom designed confinement slides. The confinement slide was polymerized in PDMS on a round 10mm standard microscope coverslip and designed with a regular holes array (diameter 449 μm, 1 mm spacing). Briefly, after activating the coverslip in a plasma chamber (Diener Electronics, Germany) for 2 min, this coverslip then was used to press a PDMS drop on top of the wafer, to obtain a thin layer. After baking at 95°C on a hot plate for 15 min, excess PDMS was removed. Isopropanol was used to peel off the glass slide with the PDMS pillars from the wafer. Microfabricated coverslips with confining pillars (8 μm height) were then attached to PDMS spacers that were stuck on a 6-well plate lid (4DCell).

### CH-STED super-resolution microscopy

For CH-STED microscopy, cells were grown as described above. Parental RPE-1 cells were seeded in the day before the experiment in coverslips coated with FBN. After fixation with 4% paraformaldehyde in cytoskeleton buffer the cells were extracted in PBS with 0.5% Triton-X100 (Sigma-Aldrich). The coverslips were incubated with the primary antibodies (rabbit anti-TPR, 1:100, NB100-2867; and mouse anti-NUPs, 1:100, Abcam 24609) in blocking solution overnight at 4°C. After washing with PBS-0.1%Tríton-X, the coverslips were incubated with the secondary antibodies (Abberior anti-rabbit STAR 580, 1:100, and STAR and Abberior anti-mouse STAR 635P, 1:100) at room temperature for 1h. Later, coverslips were washed in PBS with 0.1% Triton-X100 and sealed on a glass slide using mounting medium (20nM Tris pH 8, 0.5 N-propyl gallate, 90% glycerol).

The images were acquired with an Abberior Instruments “Expert Line” gated-STED coupled to a Nikon Ti microscope. For all the acquisitions, an oil-immersion 60x 1.4NA Plan-Apo objective (Nikon, Lambda Series) and pinhole size of 0.8 Airy units were used. The CH-STED technique creates an orthogonal direction on the STED parametric space that enables the independent tuning of both resolution and contrast using only one depletion beam in a standard STED setup (circular polarization based).

### Immunofluorescence

For the immunofluorescence experiments, cells were grown as previously described. RPE-1 parental cells were seeded in the day before the experiment in coverslips coated with FBN. After fixation with 4% Paraformaldehyde in cytoskeleton buffer (274mM NaCl, 2.2mM Na_2_HPO_4_, 10mM KCL, 0.8 mM KH_2_PO_4_, 4mM EDTA, 4mM MgCl_2_, 10mM Glucose, pH 6.1), cells were extracted with PBS-0.5% Triton-X100 (Sigma-Aldrich) following three washes (5 min each) with PBS-0.1% Triton-X100 and a 30 min incubation in blocking solution (10% FBS in 10% Triton-X100 in PBS). The coverslips were incubated with the primary antibodies (rabbit anti-cPLA2, 1:100, Cell Signalling; mouse anti-LaminA/C, 1:500, ABCAM; rat anti-tyrosinated *α*-tubulin 1:500, Bio-Rad; mouse *γ*-H2AX, 1:2000) in blocking solution for 1h at room temperature (RT). After washing with PBS-0.1%Tríton-X for 5 min, the coverslips were incubated with the secondary antibodies (Alexa Fluor 488, 568 and 647, 1:2000; Invitrogen) at RT for 1h. Later, coverslips were washed, 3x, in PBS with 0.1% Triton-X100 and 1x with PBS. Images were acquired using an AxioImager z1 (63x, Plan oil differential interference contract objective lens, 1.4 NA; from Carl Zeiss), coupled with a CCD camera (ORCA-R2; Hamamatsu Photonics) and the Zen software (Carl Zeiss).

### Quantitative image analysis

For the quantifications of cyclin B1 levels, images were analysed using ImageJ. A small square region of interest (ROI) was defined, and cyclin B1 fluorescence intensity measured, throughout time in the cell nucleus. The same ROI was used to measure the background outside the cell area. All fluorescence intensity values were then background corrected and the values were normalized to the lowest nuclear cyclin B1 level. Time zero was defined as the lowest nuclear cyclin B1 intensity inside the nucleus.

For quantification of cPLA2 fluorescence intensity on the NE, images were analysed using ImageJ. A defined ROI was used to measure the fluorescence intensity values in 5 different regions outside the cells, which was then used to calculate the average background levels. Afterwards, an equivalent ROI was used to measure fluorescence intensity in the nucleoplasm and at the NE. Images were background-subtracted and then area normalized. cPLA2 enrichment at the NE was calculated by obtaining ratio between fluorescence in the NE and in nucleoplasm.

Nuclear irregularity index (NII) was used to estimate the overall folding of the nucleus. To do so, we first obtained 2D images of the medial section of the nucleus using an anti-Lamin A/C antibody. These images were processed to obtain the nuclear area and convex area using ImageJ. Nuclear solidity was then calculated as area/convex area. Nuclear Irregularity Index was defined as 1-nuclear solidity.

### MATLAB custom algorithm for nuclear pore analysis

A computational algorithm was developed in MATLAB (The MathWorks Inc, USA; v2018b) to quantify compression-induced topological changes in the nuclear pores, within the nuclear membrane. For the analyses done in this study, we used a method focused on estimating changes on the average inter-distance between TPR and a mixture of proteins that compose the nuclear pore complex (NPC), respectively tagged with Abberior anti-rabbit STAR 580 and anti-mouse STAR 635P. Spatial periodicity on either staining (TPR and NPC) was estimated through a spatial Fast-Fourier Transform (FFT) operation on membrane cross-section images. As an alternative, and as validation, intensity profile autocorrelation was also used to assess the spatial periodicity. The entire length of the nuclear membrane was previously scanned to select only segments with low curvatures and the “Straighten” tool^47^ was used prior to the FFT/autocorrelation operations.

### MATLAB custom algorithm for centrosome tracking

To perform a detailed quantitative analysis of centrosome positioning and movement and cell rounding, we used a previously described, custom-designed MATLAB (The MathWorks Inc, USA; v2018b) script^26^. The algorithm used a specific workflow designed for centrosome tracking in a 3D space having in consideration a pixel size of 0.176 μm and a z-step of 0.5 μm. The nucleus shape was reconstructed using H2B-GFP as marker and cell shape was reconstructed using tubulin-RFP as a marker. Using this tool, we were able to correlate the angle between the centrosomes and the nucleus, as well as cell rounding during mitotic entry.

### Laser microsurgery

Laser microsurgery was performed with a doubled-frequency laser (FQ-500-532; Elforlight) coupled with an inverted microscope (TE2000U; Nikon), using a 100x 1.4NA, plan-apochromatic DIC objective lens and equipped with an iXonEM + EM-CD camera (Andor Technology). To induce a break on the NE, we used 8 consecutive pulses, with a pulse energy of 3-5 μJ and an interval of around 10ns.

### Statistical analysis

Three to six independent experiments were used for statistical analysis. Knockdown efficiency was assessed by immunoblots quantification. When data are represented as box-whisker plots, the box size represents 75% of the population and the line inside the box represents the median of the sample. The size of the bars (whiskers) represents the maximum (in the upper quartile) and the minimum (in the lower quartile) values. Statistical analysis for multiple group comparison was performed using a parametric one-way analysis of variance (ANOVA) when the samples had a normal distribution. Otherwise, multiple group comparison was done using a nonparametric ANOVA (Kruskal-Wallis). Multiple comparisons were analyzed using either post-hoc Student-Newman-Keuls (parametric) or Dunn’s (nonparametric) tests. When only two experimental groups were compared, we used either a parametric t test or a nonparametric Mann-Whitney test. Comparison for multiple time-course datasets was done using an ANOVA Repeated Measures, when the samples had a normal distribution. Otherwise, group comparison was done using Repeated Measures ANOVA on Ranks. Distribution normalities were assessed using the Kolmogorov–Smirnov test. No power calculations were used. All statistical analyses were performed using SigmaStat 3.5 (Systat Software, Inc.).

## Acknowledgments

This work was funded by Portuguese funds through FCT—Fundação para a Ciência e a Tecnologia/Ministério da Ciência, Tecnologia e Ensino Superior in the framework of the project PTDC/BIA-CEL/6740/2020. M.D. is supported by grant PD/BD/135548/2018 from the BiotechHealth FCT-funded PhD program. Work in the Maiato lab is funded by the European Research Council (ERC) consolidator grant CODECHECK, under the European Union’s Horizon 2020 research and innovation programme (grant agreement 681443), Fundação para a Ciência e a Tecnologia of Portugal (PTDC/MED-ONC/3479/2020), and the *NORTE-01-0145-FEDER-000051* project supported by NORTE 2020 under the PORTUGAL 2020 Partnership Agreement through the European Regional Development Fund. The authors would like to thank Jonathon Pines for the gift of the RPE-1 cyclin B1-Venus and HeLa cyclin B1-Venus cell lines and the plasmid for expression of cyclin B1-5A-GFP. The authors thank Dr. Buzz Baum, Dr. Alexis Lomakin and members of the Ferreira and Maiato labs for critical reading of the manuscript.

## Author contributions

M.D. performed experimental work, analyzed data, prepared figures and wrote the manuscript. J.G.F. provided the conceptual framework, analyzed data, prepared figures, and wrote the manuscript. H.M. provided access to essential equipment, reviewed, and edited the manuscript. A.O. and P.A developed MATLAB computational tools, reviewed, and edited the manuscript.

## Data availability

The data that support the findings included in this manuscript are available from the corresponding author upon reasonable request.

## Code availability

All custom-designed computational tools used in this manuscript are available from the corresponding author upon reasonable request.

---

**Supplementary Figure 1**

**(a)** Schematic representation of the dynamic confiner used to perform the experiments. **(b)** RPE-1 cell expressing Lap2β-mRFP under confinement. Please note that, upon confinement, the NE unfolds without apparent breaks. Time frame is in seconds and scale bar corresponds to 10μm. Images were acquired with a 20sec interval. **(c)** Representative images of prophase nuclei of parental RPE-1 cells stained with TPR (magenta) and a NPC mix (green) in control, non-confined conditions (top panel; n=44) and under confined conditions (bottom panel; n=55), acquired with CH-STED. Scale bar corresponds to 10μm. **(d)** Distance between neighbouring nuclear pores (NPC-NPC distance) in control, non-confined (black) and confined conditions (green) for prophase and interphase cells (*p<0.001). Measurements were done using a custom MATLAB algorithm, as described in the Materials and Methods section. **(e)** Representative immunofluorescence images of parental RPE-1 prophase cells treated with Etoposide (positive control), cells seeded on glass, seeded on glass and under confinement or cells seeded on a 5kPa hydrogel, stained for histone *γ*-H2AX and DAPI. **(f)** Quantification of the number of *γ*-H2AX foci per cell for the different treatments/seeding conditions (***p<0.001).

**Supplementary Figure 2**

RPE-1 cell expressing cyclin B1-Venus and tubulin-mRFP dividing on an FBN-coated substrate under control conditions **(a)** or subjected to a hypotonic shock **(b)**. Time frame is in seconds and scale bar corresponds to 10μm. Images were acquired with a 20sec interval and time zero corresponds to NEP. **(c)** Normalized nuclear cyclin B1 fluorescence accumulation over time in control cells (black; n=10) and in cells subjected to the hypotonic shock (green; n=19; ***p<0.001). Time zero corresponds to NEP and the dashed line represents the moment of the hypotonic shock. **(d)** HeLa cell expressing cyclin B1-Venus and H2B-mRFP dividing on an FBN-coated substrate. Time frame in seconds and scale bar corresponds to 10μm. Images were acquired with 20sec interval and time zero corresponds to the first imaged frame. The lower panel highlights nuclear accumulation of cyclin B1 in the same cell. **(e)** HeLa cell expressing cyclin B1-Venus and tubulin-mRFP dividing on an FBN-coated substrate under confinement. Time frame in seconds and scale bar corresponds to 10μm. Images were acquired with 20sec interval and time zero corresponds to the first imaged frame. The lower panel highlights the nuclear accumulation of cyclin B1 in the same cell. **(f)**: Normalized nuclear cyclin B1 fluorescence accumulation over time in control (black; n=6) and in confined cells (green; n=6). Time zero corresponds to the lowest intensity value inside the cell nucleus.

**Supplementary Figure 3**

**(a)** Normalized nuclear cyclin B1 fluorescence accumulation over time in RPE-1 control cells (black; n=10) and in RPE-1 cells expressing the Rap1* (green; n=19). Time zero corresponds to NEP. **(b)** Cell membrane eccentricity in control cells (black) and cells expressing Rap1* (green; ***p<0.001). **(c)** RPE-1 cell expressing cyclin B1-5A-GFP (top panel) and tubulin-mRFP (bottom panel) seeded on an FBN-coated substrate and under confinement (n=19). Time frame in seconds and scale bar corresponds to 10μm. Images were acquired with 20sec interval and time zero corresponds to the first imaged frame. **(d)** Nuclear accumulation of cyclinB1-5A-GFP (black) and tubulin (green), over time. Time zero corresponds to the lower intensity value inside the nucleus. The vertical dashed line corresponds to the moment of confinement. RPE-1 cells expressing cyclin B1-Venus and tubulin-mRFP seeded on an FBN-coated substrate, treated with Plk1 inhibitor (Plk1i) without **(e**; n=25) or with **(f**; n=12) confinement. Time frame in seconds and scale bar corresponds to 10μm. Images were acquired with 20sec interval and time zero corresponds to NEP. **(g)** Percentage of cells treated with Plk1i without and with mechanical confinement that enter mitosis, when compared to cells treated with DMSO. **(h)** Nuclear accumulation of cyclin B1 over time, in cells treated with DMSO (black), cells treated with Plk1i (red), and cells confined cells treated with Plk1i under confinement that either enter (green) or fail to enter (magenta) mitosis (***p<0.001). Time zero corresponds to the lowest fluorescence intensity value of cyclin B1 inside the nucleus

**Supplementary Figure 4**

**(a)** Representative immunofluorescence images of parental RPE-1 cells in prophase with or without confinement, after treatment with Y-27632 (n=29) or p-N-blebb (n=25), stained for cPLA2, DAPI and Lamin A/C. Scale bars, 10μm. **(b)** Distance between neighbouring nuclear pore complexes (NPC-NPC distance) in cells treated with p-N-blebb without (black) or with confinement (green; ***p<0.001). This measurement was done using the custom MATLAB algorithm, as described in the Materials and Methods section. **(c)** Representative images of parental RPE-1 cells stained for TPR (magenta) and an NPC mix (green) in non-confined (top panel; n=62) and confined conditions (bottom panel; n=59), after treatment with p-N-blebb, acquired with CH-STED. Scale bar corresponds to 10μm. **(d)** RPE-1 cell expressing cyclin B1-Venus and tubulin-mRFP dividing on an FBN-coated substrate treated with BAPTA-AM + 2APB in non-confined conditions (top panel; n=17) and confined conditions (lower panel; n=11). Time frame in seconds and scale bar corresponds to 10μm. Images were acquired with 20sec interval and time zero corresponds to NEP. **(e)** RPE-1 cell expressing cyclinB1-Venus and tubulin-mRFP dividing on an FBN-coated substrate treated with AACOCF3 in non-confined conditions (top panel; n=12) and confined conditions (lower panel; n=15). Time frame in seconds and scale bar corresponds to 10μm. Images were acquired with 20sec interval and time zero corresponds to NEP.

**Supplementary Figure 5**

**(a)** RPE-1 cell expressing H2B-GFP and tubulin-mRFP dividing on an FBN-coated substrate treated with STLC (Eg5 inhibitor) without (top panel; n=18) and with confinement (lower panel; n=26). Time frame in min:sec and scale bar corresponds to 10μm. Images were acquired with a 20sec interval and time zero corresponds to NEP. **(b)** Pole-to-pole distance on STLC treated and non-confined cells (black) and STLC treated and confined cells (green; n.s. – not significant). **(c)** Representative image of a western blot showing the depletion of DHC with RNAi. Vinculin was used as loading control. **(d)** RPE-1 cell expressing H2B-GFP and tubulin-mRFP dividing on an FBN-coated substrate treated with DHC RNAi without (top panel; n=19) and with confinement (lower panel; n=19). Time frame in min:sec and scale bars correspond to 10μm. Images were acquired with a 20sec interval and time zero corresponds to NEP. **(e)** Distribution of centrosome angles relative to the nucleus (measured at the moment of NEP) in cells with DHC RNAi treatment and without (top panel) or with confinement (bottom panel).

## References

1. Lancaster, O. et al. Mitotic Rounding Alters Cell Geometry to Ensure Efficient Bipolar Spindle Formation. Dev. Cell 25, 270–283 (2013).

2. Cattin, C. J. et al. Mechanical control of mitotic progression in single animal cells. Proc. Natl. Acad. Sci. U. S. A. 112, 1502029112– (2015).

3. Gavet, O. & Pines, J. Progressive Activation of CyclinB1-Cdk1 Coordinates Entry to Mitosis. Dev. Cell (2010). doi:10.1016/j.devcel.2010.02.013

4. Dantas, M., Lima, J. T. & Ferreira, J. G. Nucleus-Cytoskeleton Crosstalk During Mitotic Entry. 9, 1–9 (2021).

5. Vitiello, E. et al. Acto-myosin force organization modulates centriole separation and PLK4 recruitment to ensure centriole fidelity. Nat. Commun. (2019). doi:10.1038/s41467-018-07965-6

6. Uroz, M. et al. Regulation of cell cycle progression by cell–cell and cell–matrix forces. Nat. Cell Biol. 1 (2018). doi:10.1038/s41556-018-0107-2

7. Lindqvist, A., Rodríguez-Bravo, V. & Medema, R. H. The decision to enter mitosis: feedback and redundancy in the mitotic entry network. Journal of Cell Biology (2009). doi:10.1083/jcb.200812045

8. Hagting, A., Karlsson, C., Clute, P., Jackman, M. & Pines, J. MPF localization is controlled by nuclear export. EMBO J. (1998). doi:10.1093/emboj/17.14.4127

9. Toyoshima, F., Moriguchi, T., Wada, A., Fukuda, M. & Nishida, E. Nuclear export of cyclin B1 and its possible role in the DNA damage-induced G2 checkpoint. EMBO J. (1998). doi:10.1093/emboj/17.10.2728

10. Pines, J. & Hunter, T. Isolation of a human cyclin cDNA: Evidence for cyclin mRNA and protein regulation in the cell cycle and for interaction with p34cdc2. Cell (1989). doi:10.1016/0092-8674(89)90936-7

11. Abe, S. et al. The initial phase of chromosome condensation requires Cdk1-mediated phosphorylation of the CAP-D3 subunit of condensin II. Genes Dev. (2011). doi:10.1101/gad.2016411

12. Huang, S., Chen, C. S. & Ingber, D. E. Control of cyclin D1, p27(Kip1), and cell cycle progression in human capillary endothelial cells by cell shape and cytoskeletal tension. Mol Biol Cell (1998).

13. Gudipaty, S. A. et al. Mechanical stretch triggers rapid epithelial cell division through Piezo1. Nature (2017). doi:10.1038/nature21407

14. Benham-Pyle, B. W., Pruitt, B. L. & Nelson, W. J. Mechanical strain induces E-cadherin-dependent Yap1 and β-catenin activation to drive cell cycle entry. Science (80-.). (2015). doi:10.1126/science.aaa4559

15. Klein, E. A. et al. Cell-Cycle Control by Physiological Matrix Elasticity and In Vivo Tissue Stiffening. Curr. Biol. (2009). doi:10.1016/j.cub.2009.07.069

16. Aureille, J. et al. Nuclear envelope deformation controls cell cycle progression in response to mechanical force. EMBO Rep. (2019). doi:10.15252/embr.201948084

17. Vianay, B. et al. Variation in traction forces during cell cycle progression. Biol. Cell 110, 91–96 (2018).

18. Lombardi, M. L. & Lammerding, J. Keeping the LINC: The importance of nucleocytoskeletal coupling in intracellular force transmission and cellular function. in Biochemical Society Transactions (2011). doi:10.1042/BST20110686

19. Arsenovic, P. T. et al. Nesprin-2G, a Component of the Nuclear LINC Complex, Is Subject to Myosin-Dependent Tension. Biophys. J. (2016). doi:10.1016/j.bpj.2015.11.014

20. Elosegui-Artola, A. et al. Force Triggers YAP Nuclear Entry by Regulating Transport across Nuclear Pores. Cell (2017). doi:10.1016/j.cell.2017.10.008

21. Lomakin, A. J. et al. The nucleus acts as a ruler tailoring cell responses to spatial constraints. Science (80-.). (2020). doi:10.1126/science.aba2894

22. Jacchetti, E. et al. The nuclear import of the transcription factor MyoD is reduced in mesenchymal stem cells grown in a 3D micro-engineered niche. Sci. Rep. (2021). doi:10.1038/s41598-021-81920-2

23. Nava, M. M. et al. Heterochromatin-Driven Nuclear Softening Protects the Genome against Mechanical Stress-Induced Damage. Cell (2020). doi:10.1016/j.cell.2020.03.052

24. Swift, J. et al. Nuclear lamin-A scales with tissue stiffness and enhances matrix-directed differentiation. Science (80-.). (2013). doi:10.1126/science.1240104

25. Venturini, V. et al. The nucleus measures shape changes for cellular proprioception to control dynamic cell behavior. Science (80-.). (2020). doi:10.1126/science.aba2644

26. Nunes, V. et al. Centrosome–nuclear axis repositioning drives the assembly of a bipolar spindle scaffold to ensure mitotic fidelity. Mol. Biol. Cell (2020). doi:10.1091/mbc.e20-01-0047

27. Le Berre, M., Zlotek-Zlotkiewicz, E., Bonazzi, D., Lautenschlaeger, F. & Piel, M. Methods for two-dimensional cell confinement. Methods Cell Biol. 121, 213–229 (2014).

28. Bakkenist, C. J. & Kastan, M. B. DNA damage activates ATM through intermolecular autophosphorylation and dimer dissociation. Nature (2003). doi:10.1038/nature01368

29. Kumar, A. et al. ATR mediates a checkpoint at the nuclear envelope in response to mechanical stress. Cell (2014). doi:10.1016/j.cell.2014.05.046

30. Maddox, A. S. & Burridge, K. RhoA is required for cortical retraction and rigidity during mitotic cell rounding. J. Cell Biol. (2003). doi:10.1083/jcb.200207130

31. Peter, M., Nakagawa, J., Dorée, M., Labbé, J. C. & Nigg, E. A. In vitro disassembly of the nuclear lamina and M phase-specific phosphorylation of lamins by cdc2 kinase. Cell (1990). doi:10.1016/0092-8674(90)90471-P

32. Heald, R. & McKeon, F. Mutations of phosphorylation sites in lamin A that prevent nuclear lamina disassembly in mitosis. Cell (1990). doi:10.1016/0092-8674(90)90470-Y

33. Yang, J., Bogerd, H. P., Wang, P. J., Page, D. C. & Cullen, B. R. Two closely related human nuclear export factors utilize entirely distinct export pathways. Mol. Cell (2001). doi:10.1016/S1097-2765(01)00303-3

34. Moore, J. D., Yang, J., Truant, R. & Kornbluth, S. Nuclear import of Cdk/cyclin complexes: Identification of distinct mechanisms for import of Cdk2/cyclin E and Cdc2/cyclin B1. J. Cell Biol. (1999). doi:10.1083/jcb.144.2.213

35. Hagting, A., Jackman, M., Simpson, K. & Pines, J. Translocation of cyclin B1 to the nucleus at prophase requires a phosphorylation-dependent nuclear import signal. Curr. Biol. (1999). doi:10.1016/S0960-9822(99)80308-X

36. Li, J., Meyer, A. N. & Donoghue, D. J. Nuclear localization of cyclin B1 mediates its biological activity and is regulated by phosphorylation. Proc. Natl. Acad. Sci. U. S. A. (1997). doi:10.1073/pnas.94.2.502

37. Santos, S. D. M., Wollman, R., Meyer, T. & Ferrell, J. E. Spatial positive feedback at the onset of mitosis. Cell (2012). doi:10.1016/j.cell.2012.05.028

38. Gheghiani, L., Loew, D., Lombard, B., Mansfeld, J. & Gavet, O. PLK1 Activation in Late G2 Sets Up Commitment to Mitosis. Cell Rep. (2017). doi:10.1016/j.celrep.2017.05.031

39. Enyedi, B., Jelcic, M. & Niethammer, P. The Cell Nucleus Serves as a Mechanotransducer of Tissue Damage-Induced Inflammation. Cell 165, 1160–1170 (2016).

40. Kao, J. P. Y., Alderton, J. M., Tsien, R. Y. & Steinhardt, R. A. Active involvement of Ca2+ in mitotic progression of Swiss 3T3 fibroblasts. J. Cell Biol. (1990). doi:10.1083/jcb.111.1.183

41. Strauss, B. et al. Cyclin B1 is essential for mitosis in mouse embryos, and its nuclear export sets the time for mitosis. J. Cell Biol. (2018). doi:10.1083/jcb.201612147

42. Furuno, N., Elzen, N. Den & Pines, J. Human cyclin A is required for mitosis until mid prophase. J. Cell Biol. (1999). doi:10.1083/jcb.147.2.295

43. Tse, H. T. K., Weaver, W. M. C. & Carlo, D. Increased asymmetric and multi-daughter cell division in mechanically confined microenvironments. PLoS One (2012). doi:10.1371/journal.pone.0038986

44. Silkworth, W. T., Nardi, I. K., Paul, R., Mogilner, A. & Cimini, D. Timing of centrosome separation is important for accurate chromosome segregation. Mol. Biol. Cell (2012). doi:10.1091/mbc.E11-02-0095

45. Schweizer, N., Pawar, N., Weiss, M. & Maiato, H. An organelle-exclusion envelope assists mitosis and underlies distinct molecular crowding in the spindle region. J. Cell Biol. (2015). doi:10.1083/jcb.201506107

46. Zimmerli, C. E. et al. Nuclear pores dilate and constrict in cellulo. Science (80-.). 374, (2021).

47. Schindelin, J. et al. Fiji: An open-source platform for biological-image analysis. Nature Methods (2012). doi:10.1038/nmeth.2019

