## Supplementary figures and images for "Mechanosensitive control of mitotic entry"

### Supplemental Figure 1

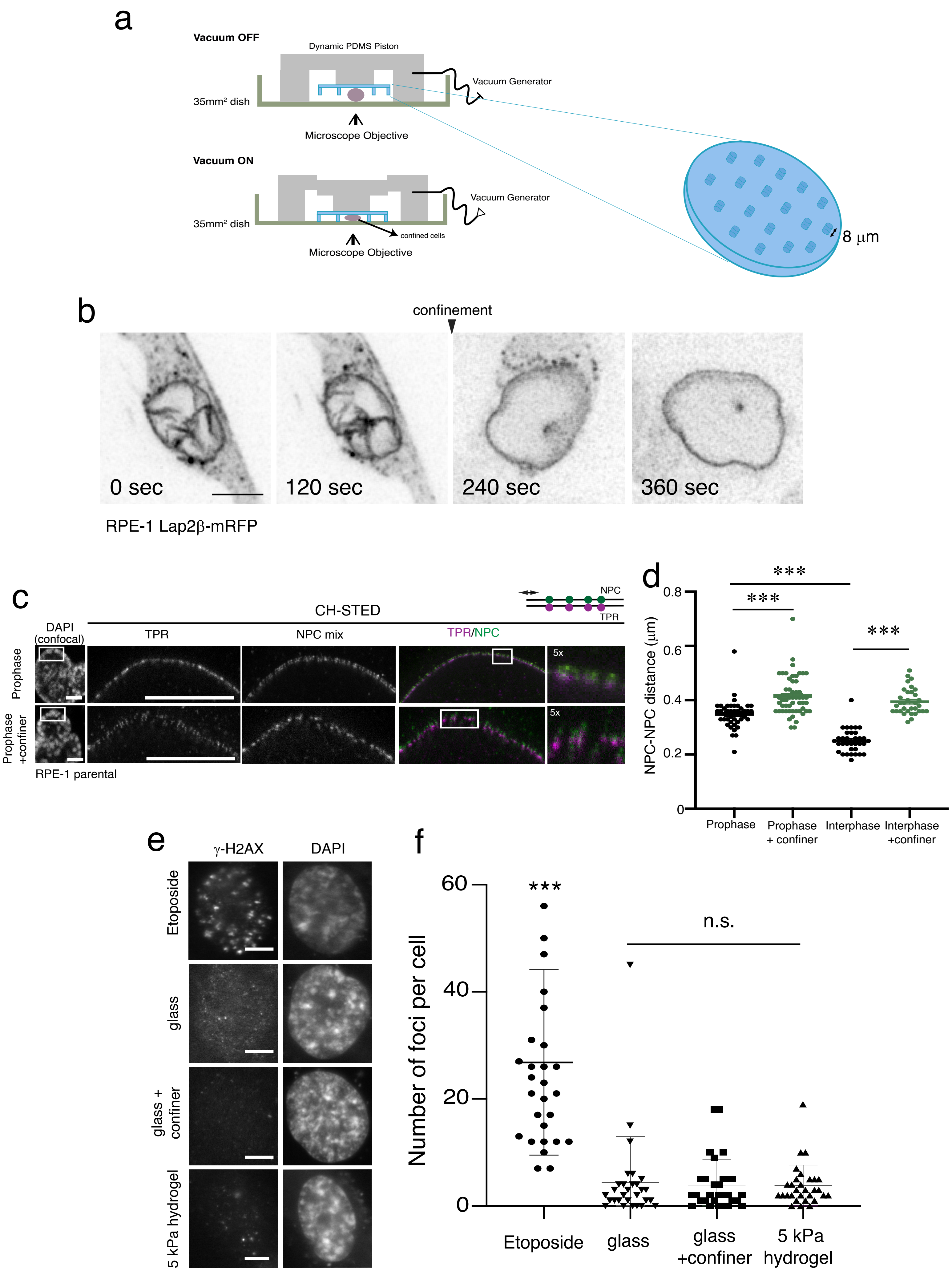


Supplementary Figure 1

### Supplemental Figure 2

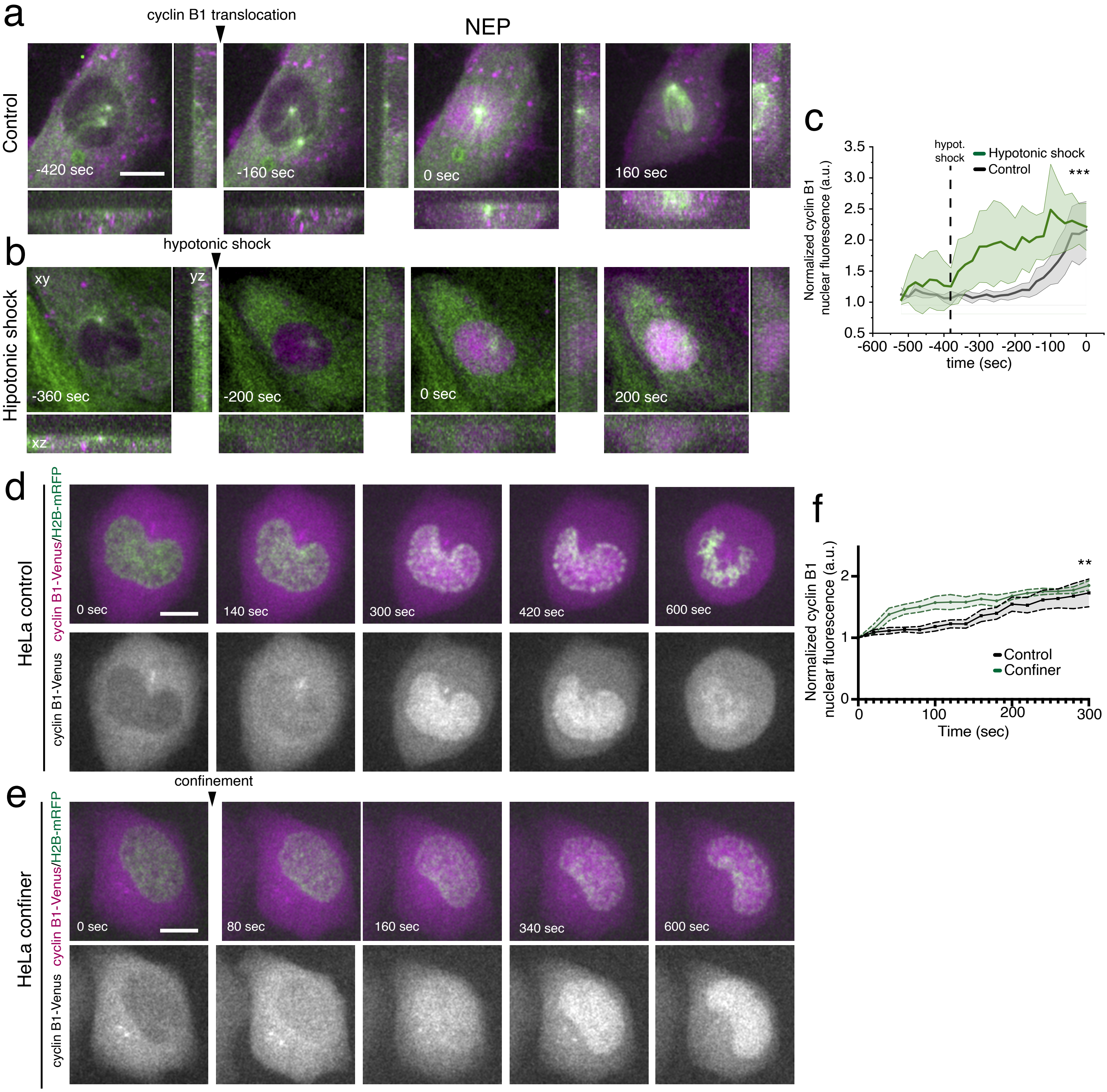


#### Supplementary Figure 2

### Supplemental Figure 3

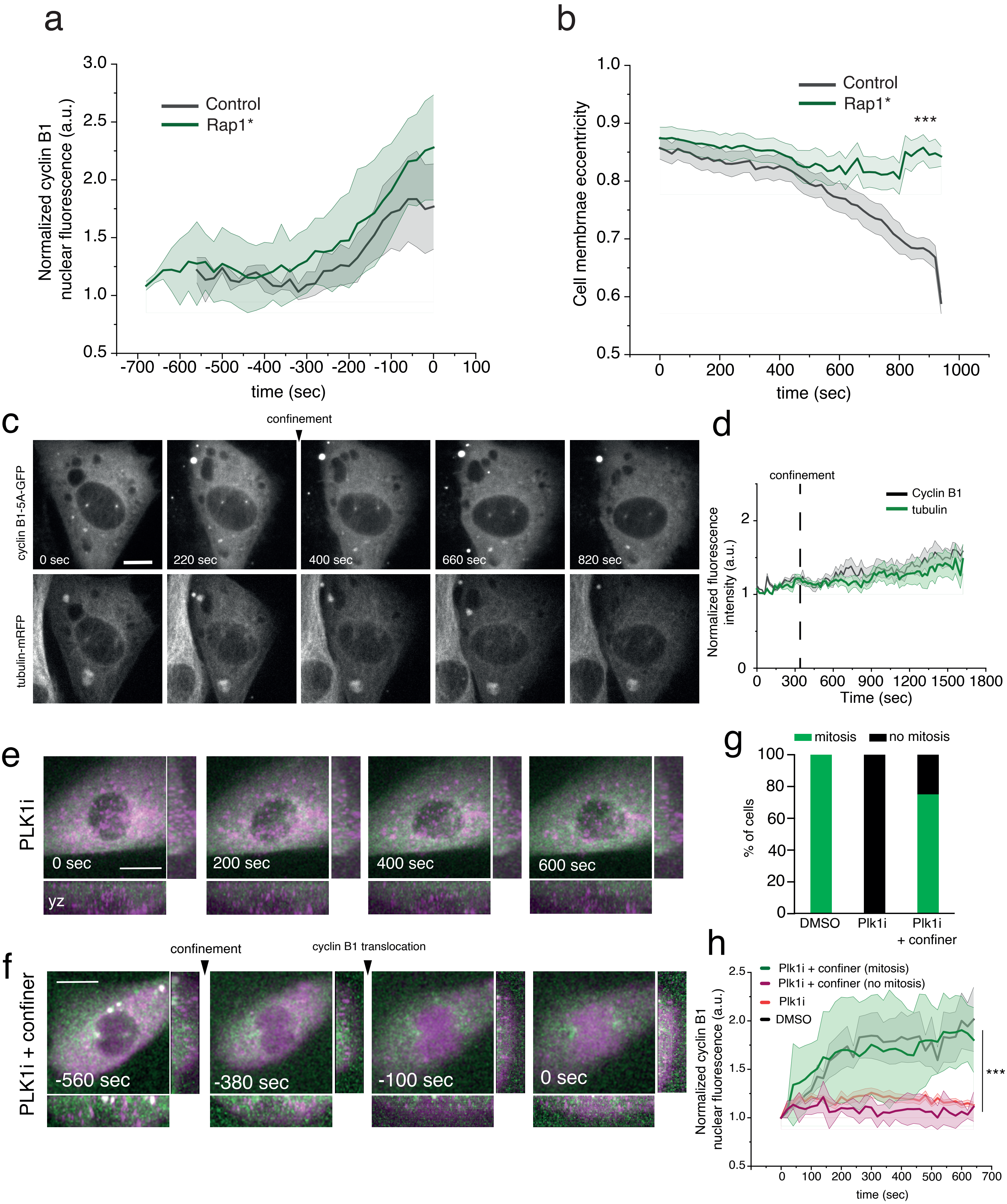


Supplementary Figure 3

### Supplemental Figure 4

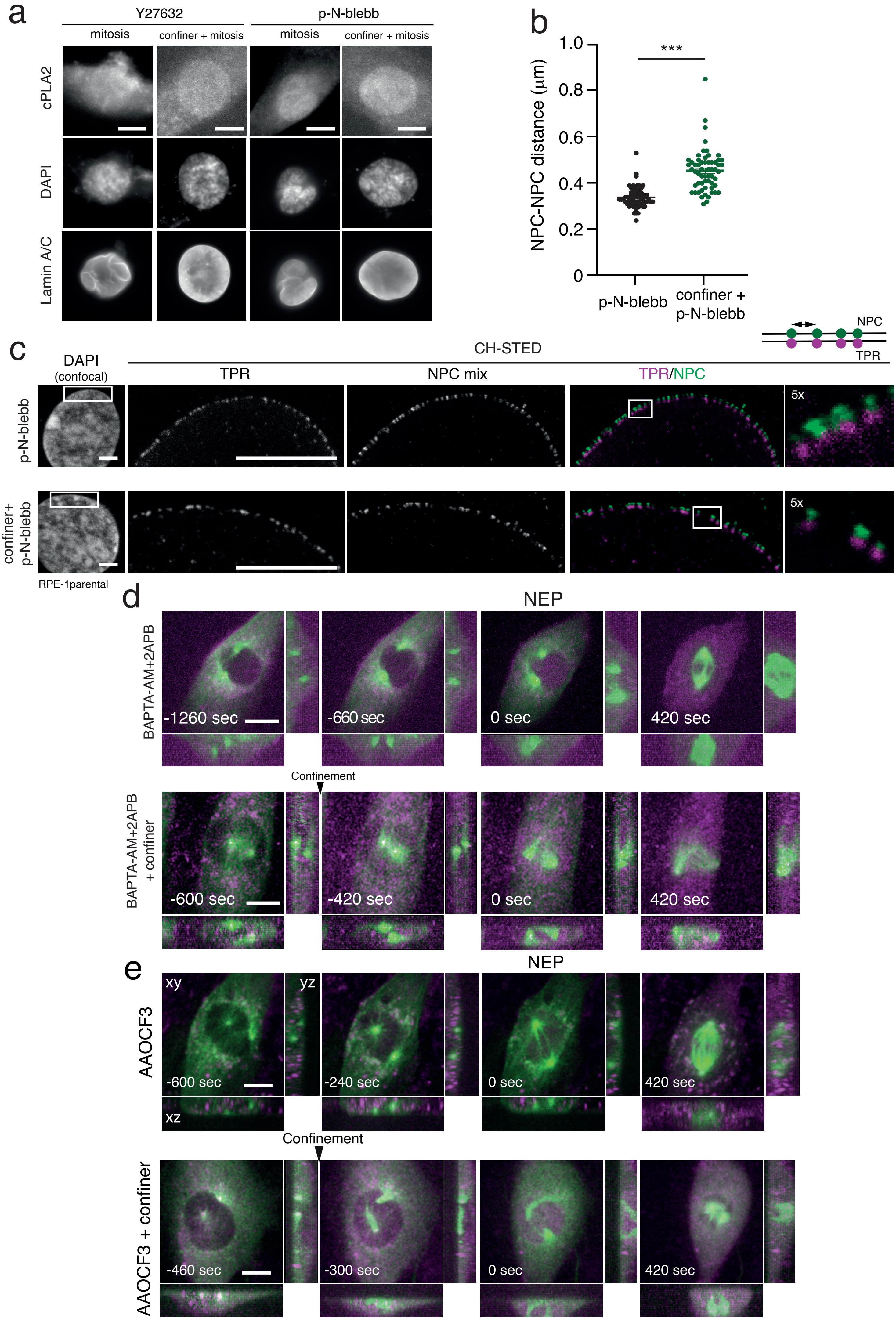


#### Supplementary Figure 4

### Supplemental Figure 5

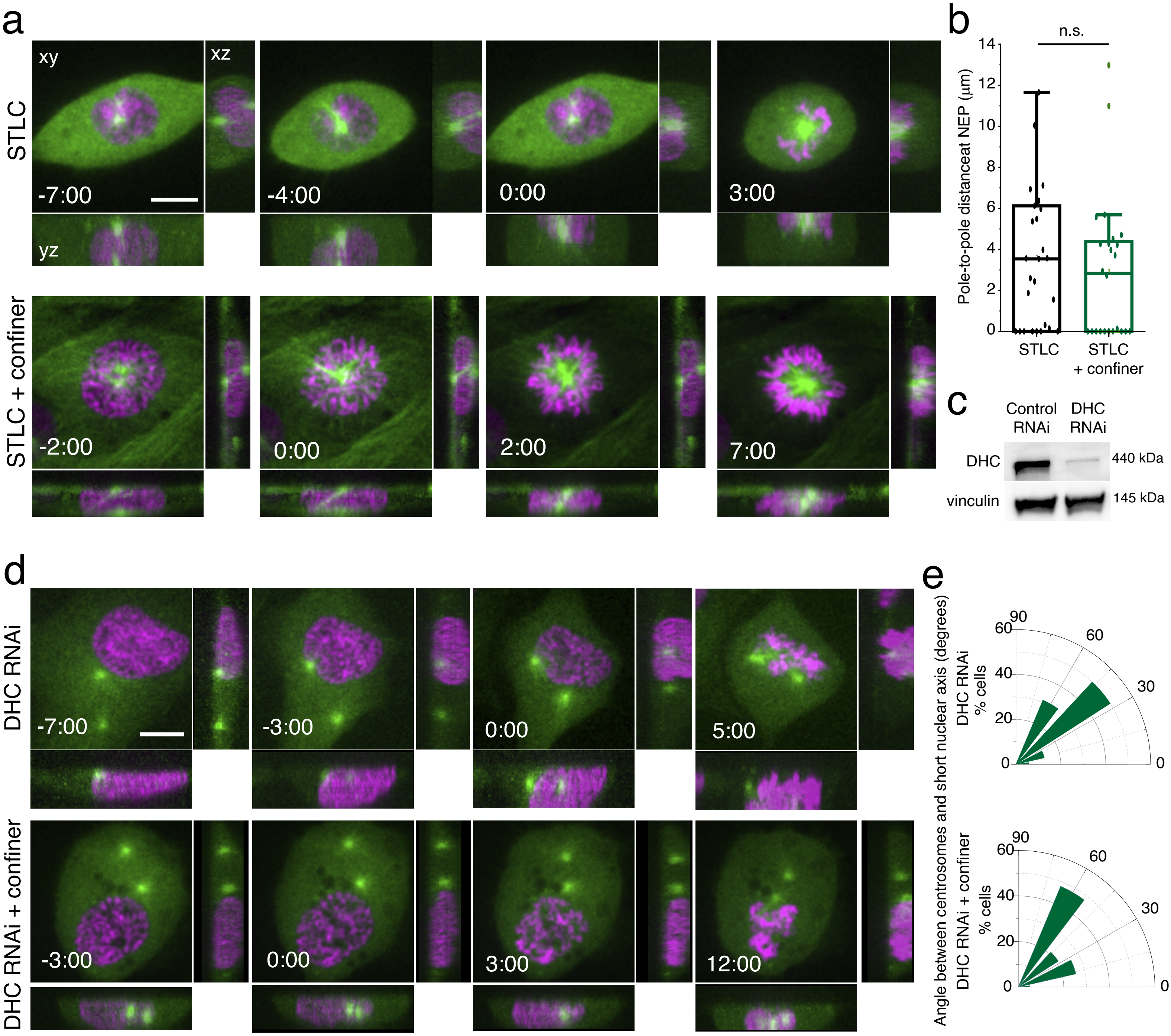


Supplementary Figure 5
